# Genomic local adaptation of a generalist plant species to pollinator communities and abiotic factors

**DOI:** 10.1101/2022.08.05.502924

**Authors:** L. Frachon, L. Arrigo, Q. Rusman, L. Poveda, W. Qi, G. Scopece, F.P. Schiestl

## Abstract

The reproductive success of generalist flowering plants is influenced by a complex ecological network that includes interactions with a diverse pollinator community and abiotic factors. However, knowledge about of the adaptative potential of plants to complex ecological networks and the underlying genetic mechanisms is still limited. Based on a pool-sequencing approach of 21 natural populations of *Brassica incana* in Southern Italy, we combined a genome-environmental association analysis with a genome scan for signature of selection to discover genetic variants associated with ecological variation. We demonstrated that *B. incana* is locally adapted both to the identity of functional categories and overall pollinator interactions. Interestingly, we observed only few shared candidate genes associated with long-tongue bees, soil texture, and temperature variation. Our results highlight the genomic architecture of generalist flowering plant adaptation to complex biotic interactions, and the importance of considering multiple environmental factors to describe the adaptive landscape of plant populations.

## Introduction

In natural populations, most of the flowering plant species interact simultaneously with different functional groups of pollinators (called generalist species) ensuring their reproductive success (Albrecht et al. 2012). By interacting with an assemblage of generalist and specialist pollinators, widely distributed generalist plant species (Waser et al. 1996, Johnson and Steiner 2000) appear robust to pollinator changes within mutualistic networks (Bascompte and Jordano 2007, Thébault and Fontaine 2010, Burkle et al. 2013, Zografou et al 2021). Although generalist plant species are keystones in mutualist interaction networks, we know little about the adaptive potential of these plants to pollinator communities. Only a handful of studies have demonstrated the role of pollinator assemblages on floral evolution in generalist plant species (Gomez et al. 2009, Sahli and Conner 2011, Gomez et al. 2015, Sobral et al. 2015, Schiestl et al. 2018, de Manincor et al. 2021). For instance, it has been recently showed that pollinator communities can drive flower shape evolution in generalist species *Erysimum* (Gomez et al. 2015), or geographic variation in flower scent (de Manincor et al. 2021). However, floral evolution in generalist plant species appears to be complex (Gomez et al. 2015), probably involving independent and genetically linked phenotypic traits associated with pollinator preferences (Frachon et al. 2021, Ohashi et al 2021). To understand if and how generalist plant species locally adapt to their pollinator communities, we need to investigate the underlying genomics. This will help us understand the coevolution of generalist plant-pollinator networks.

Plant-pollinator interactions are influenced by abiotic factors (Tylianakis et al. 2008, Chamberlain et al 2014, Antiqueira et al. 2020). For instance, climate change can induce mismatches between plant and pollinators due to non-synchronized phenology shifts (Hegland et al. 2009, Petanidou et al. 2014), or changes in plant attractiveness to pollinators (Petanidou and Smets 1996, Hoover et al. 2012, Herrera and Medrano 2017, Descamps et al. 2021). Moreover, soil heterogeneity can strongly affect plant attractiveness to pollinators through changes in nectar secretion, production of pollen or essential oils, and floral scent (Burkle and Irwin 2009, Majetic et al. 2017, David et al. 2019, Carvalheiro et al. 2021). Understanding how both pollinator communities and abiotic factors simultaneously drive the evolution of combined phenotypic traits and their associated genomic regions is a current challenge requiring a holistic approach from ecology to genomics (Clare et al. 2013, Lopez-Goldar and Agrawal 2021).

Genome-environment association (GEA) analysis is a powerful approach to identify genomic regions involved in the adaptive response of organisms to complex ecological networks without phenotypic characterization (de Mita et al. 2013). This approach takes advantage of the genetic fingerprint left by selective pressures due to environmental variation among natural populations. Although commonly used to understand the genetic architecture of plants involved in responses to climate change (Hancock et al. 2011, Lasky et al. 2015, Pluess et al. 2016, Cortés and Blair 2018, Frachon et al. 2018), the GEA approach has recently shown its effectiveness in unravelling the genetic variants of *A. thaliana* underlying adaptative response to complex biotic interactions such as leaf microbiome (Horton et al. 2014) and plant communities (Frachon et al. 2019).

In our study, we adopted a GEA approach to understand the adaptative potential of the generalist plant *Brassica incana* to its pollinator community (visitation by pollinator functional categories, and plant-pollinator interactions indices) as well as to potential interacting effects with climatic and edaphic (soil composition and texture) variables. By characterizing 61 ecological factors, *de novo* assembly of the *B. incana* reference genome, and pool-sequencing of 21 natural populations of *B. incana* in Southern Italy for 5’530’708 Single Nucleotide Polymorphisms (SNPs), we finely mapped QTLs associated with variation in pollinator communities, climate, and soil. This approach was combined with a genome scan for signatures of selection and enrichment in SNPs with high genetic differentiation to detect signatures of selection. Altogether, we found local adaptation of generalist plant species to a complex ecological network underlying variable genetic architecture.

## Results

### Variation in ecological variables among 21 natural populations of *Brassica incana*

Pollinator communities were characterized during the spring seasons of 2018 and 2019 by observing pollinator visitation to plants of 21 natural populations of *B. incana* (**Fig. 1, Table S1**). Flower visitors were grouped into 12 functional categories *ie* bumblebees, long-tongue bees, other large bees (called large bees), small bees, honeybees, large wasps, small flies, large flies, hoverflies, small beetles, large beetles, butterflies (**Fig. 2**). To characterize differences in pollinator communities among populations, we performed a *B. incana* – pollinators’ interaction analysis based on the total number of pollinator visits by functional categories. Pollinator communities were mainly dominated by long-tongue bees, small bees, honeybees, large bees, hoverflies, and bumblebees in decreasing order (**Fig. 2**). Moreover, the visits of functional categories of pollinators varied among 21 populations (**Fig. 2**), leading to variation in α-diversity, estimated by Shannon index, ranking from 0.63 to 1.80 (average = 1.28) (**Table S2**). The calculated indexes from the interaction analysis (**Table S2**) showed that *B. incana* plants in natural populations relied on a large number of functional categories of pollinators as suggested by the low value of species strength (minimum = 0.03, maximum = 1.71, average = 0.57, **Table S2**). Moreover, other indices confirmed that *B. incana* was a generalist plant species such as a high normalised degree index showing a high number of realized *B. incana* — pollinators links among populations (minimum = 0.17, maximum = 0.83, average = 0.56, **Table S2**) and low d-index values (minimum = 0.04, maximum = 0.53, average = 0.17, **Table S2**).

**Figure 1.**
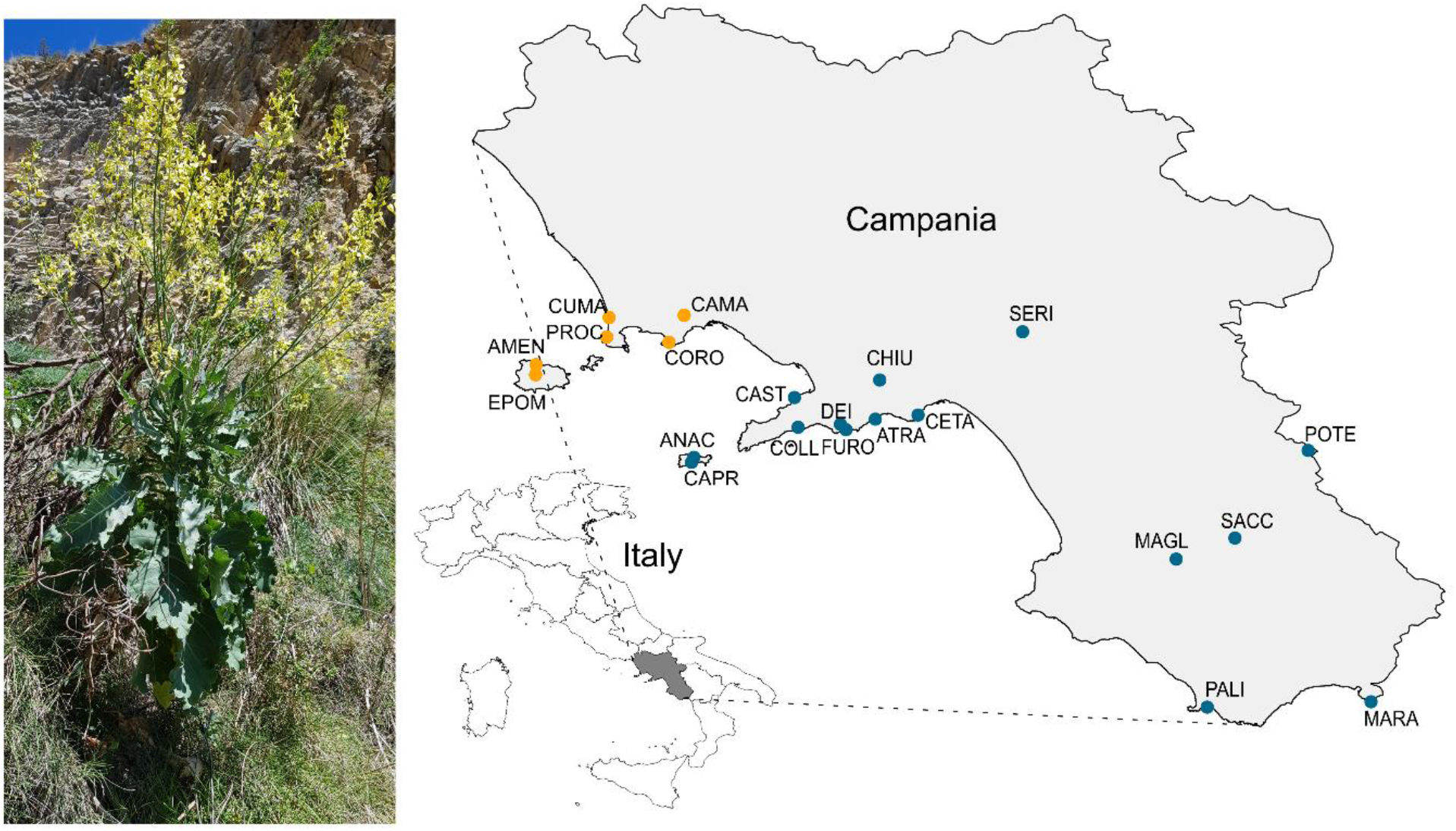
Distribution of *Brassica incana* natural populations. On the left, a photograph of a flowering *B. incana* in the PALI population. On the right, the Campania region is represented in dark grey on the Italy map. The 21 natural populations of *B. incana* are indicated with coloured dots on the map. The orange dots indicate six populations on tuff soil, and the blue dots 15 populations on limestone soil.

**Figure 2.**
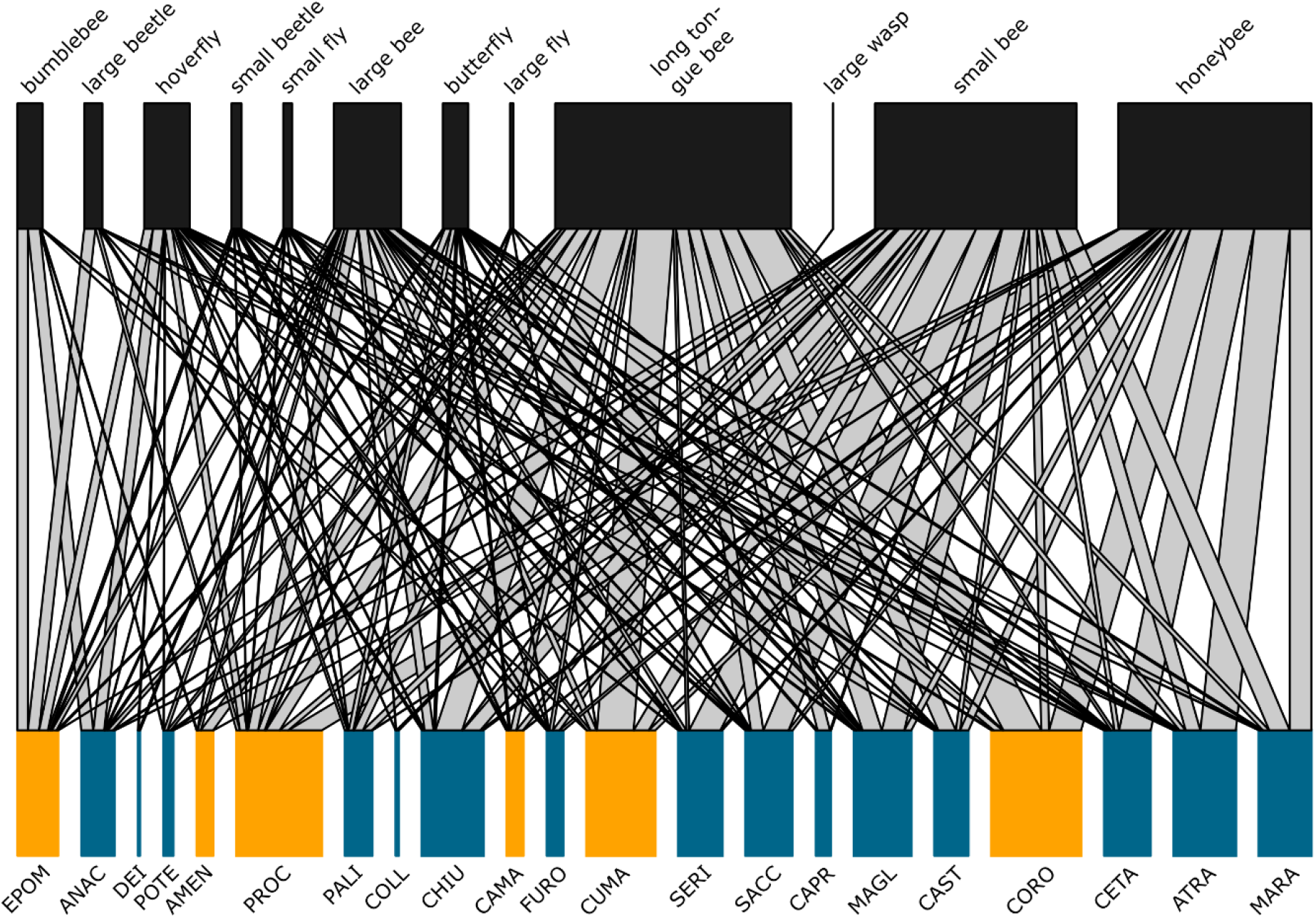
*Brassica incana* – functional categories of pollinators interaction analysis in 21 natural populations in springs 2018 and 2019. The upper part of the figure represents the 12 functional categories of pollinators. The size of boxes represents the total number of visits per functional category of pollinators observed in all 21 populations combined. The lower part the figure represents the 21 natural populations of *B. incana* coloured according to their soil type (tuff soil in orange, limestone soil in blue). The size of the boxes represents the total number of visits, all categories of pollinators combined, per population. The width of the lines connecting functional categories of pollinators to populations indicates the proportion of visits observed per pollinator category within each population.

Among the 21 natural populations, 28 of 61 characterized ecological variables were highly correlated (Spearman rho > 0.8) and were all discarded from the genomic analyses (**Fig. S1**). These highly correlated variables concerned mainly *B. incana* – pollinator interaction (four out of eight variables) and climate variables (17 variables out of 20). While we had a clear differentiation of tuff vs limestone soils following the Northwest – Southeast axis among the 21 populations (due to the geography of Southern Italy), the different characteristics of these soils were not correlated among each other’s (**Fig. S1**). The principal component analysis showed that the six populations in tuff soil (AMEN, CAMA, CORO, CUMA, PROC, EPOM) were ecologically similar and differentiated from other populations by the visits of hoverflies, the visits of large beetles, the mean annual precipitation, and the species strength (an index for plant-pollinator interactions, **Fig. S2**). The visits of bumblebees, large bees, and d index an index of plant-pollinator interactions) were correlated to the fine sand, coarse silt, Fe and summer precipitation in the ecological space created by the two first axis of the PCA (**Fig. S2**).

### Annotated reference genome of *Brassica incana*

The final assembled sequences of the *Brassica incana* reference genome were organised into 1’339 contigs, which were scaffolded into 139 super-scaffolds using Bionano optical map (**Table S5**). The 139 super-scaffolds were used in our study, including a total sequence length of 617Mbp, scaffold N50 of 12 Mbp and a longest sequence at 32 Mbp, with a BUSCO completeness score of 97.7% (**Table S5**). Sequencing data from Pacbio and Illumina used for this study are available at the European Nucleotide Archive ENA database (accession number PRJEB54646). The bionano raw data and assembled optical maps are available at National Library for Biotechnology Information (NCBI) database (sample name PRJNA859008).

In total 51’001 genes were predicted, including 50’895 proteins (from the iprscan) divided into 1’112 different categories of GO terms. As comparison, the reference genome of *Brassica oleracea* (genome size = 488.6 Mb) was composed of 53’125 genes, and *Arabidopsis thaliana* 38’311 genes (genome size = 119.1 Mb) in the NCBI database.

### Genomic architecture of *B. incana* response to ecological variations

After mapping the 21 pool-sequences from 21 natural populations to the *B. incana* reference genome we generated, we estimated the allele-frequencies across the 139 super-scaffolds for a final number of 5’530’708 SNPs. Using singular value decomposition (SVD) of the population variance-covariance matrix Ω, we estimated a strong degree of subdivision (*i*.*e*. “structuration” of populations without genetically similar populations) among our populations represented by the first PC_genomic_ explaining 94.3% of genomic variance (**Fig. S3**). This finding was supported by a weak geographic pattern along the Northeast – Southwest axis (linear model for PC1_genomic_; latitude: *t value* = 3.24, *P* = 0.005, longitude: *t value* = 3.31, *P* = 0.004, latitude*longitude: *t value* = -3.33, *P* = 0.004, adjusted R^2^ = 47.1%). Thus, the regional geographic scale applied in our study prevents the confounding effect of population structure on the genome-environmental analysis (Frachon et al. 2018, Frachon et al. 2019). The variation of most ecological factors was weakly (non-significant) correlated with the genomic variation (**Table S6**) suggesting true positives in the genome-environmental analysis (Frachon et al. 2018, Frachon et al. 2019). However, as expected with the Northeast-Southwest axis of soil, we observed significant correlation between PC1_genomic_ and six environmental factors including type of soil (long tongue bees, species strength, ratio C/N, Fine Silt, and Zn, **Table S6**). The genome-environmental association analysis performed in our study, could underestimate genomic regions involved in these six environmental variables due to confounding effects between population structure and variation of these variables, leading to potential false negatives.

To identify the adaptive genetic loci associated with functional categories of pollinators, *B. incana* — pollinator interaction indices, climate and soil composition and texture variation, we performed a genome-wide scan for associations between standardized allele frequency variation along the 139 super-scaffolds of *B. incana* genome and 33 less correlated ecological variables using a Bayesian hierarchical model. The association scores (Bayes factors, BFdB) between the variation of genomic region and ecological variables were estimated, and a local score method was applied correcting for linkage disequilibrium. Using this method, we observed neat and narrow peaks of association across the 139 super-scaffolds for the considered ecological variables (**Fig. 3, Fig. S4**). Most of the genomic regions involved in response of *B. incana* to the variation of functional categories of pollinators were unique, except for the visits of bumblebees, hoverflies, and long-tongue bees (**Fig. S5, Fig. S6**). For instance, 56% of SNPS with the 0.05% of highest association score were uniquely associated with long-tongue bees (1’435/2’541 SNPs), 95% were uniquely associated with large bees (2’414/2’541 SNPs), and 97% were uniquely associated with honeybees (2’471/2’541 SNPs, **Fig. S5**). However, only 15% of SNPs with the highest association score were uniquely associated with bumblebee or hoverfly visits (**Fig. S5**). The latter share 22% of their SNPs with highest association score between them, and an important part of SNPs with the texture of the soil (fine silt and coarse sand, **Fig. S5**). Overall, 88% of 0.05% of SNPs with the highest association score were uniquely associated with pollinator functional categories indicated an important part of SNPs involved in *B. incana* adaptation to functional categories of pollinators (**Fig. S6**).

**Figure 3:**
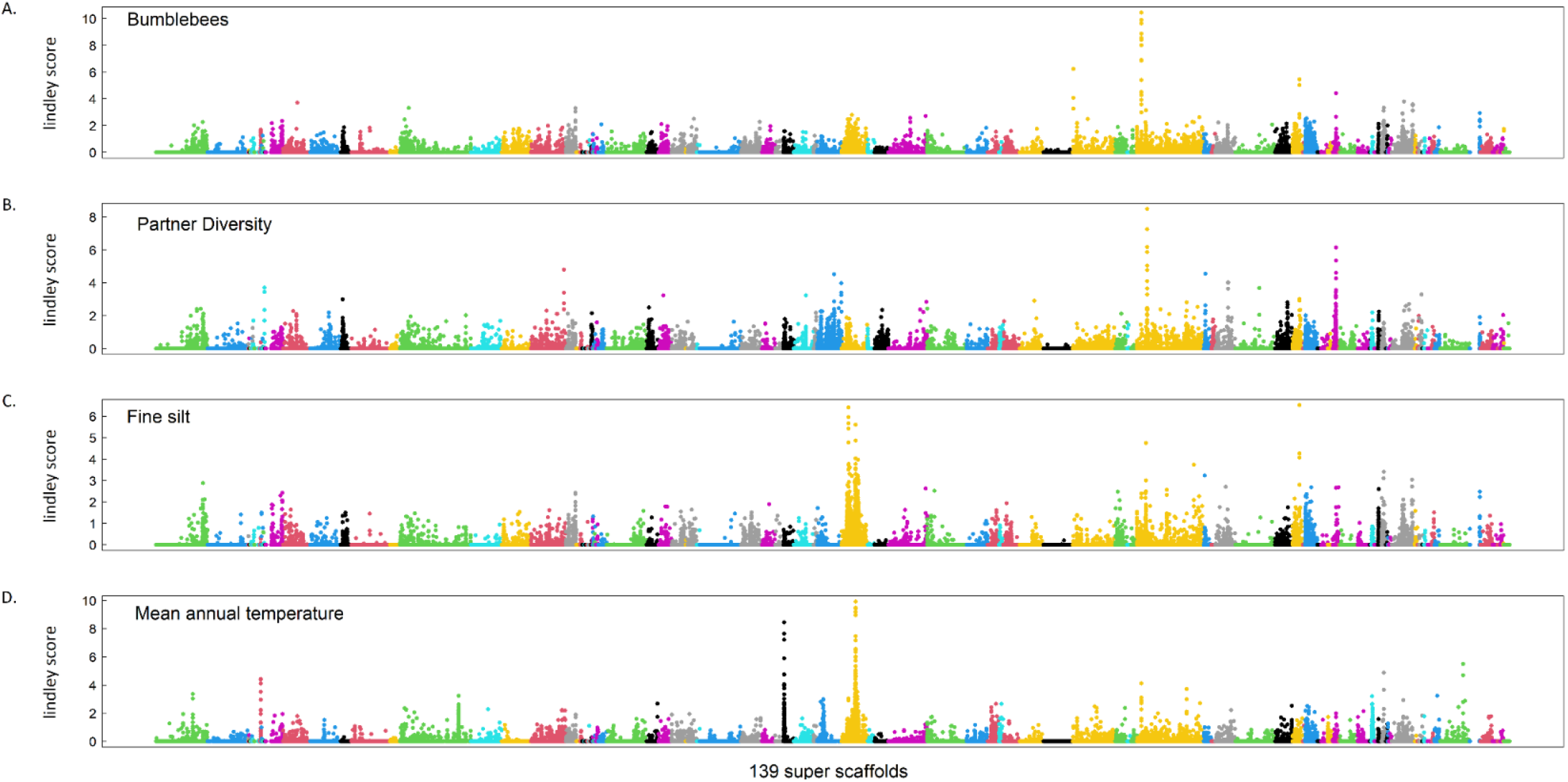
Manhattan plot of genome-environmental association analysis for four ecological variables; **(A)** visit of bumblebees, **(B)** partner diversity, **(C)** fine silt, and **(D)** mean annual temperature. The *x-axis* indicates the physical position of the 5’530’708 SNPs along the 139 super-scaffolds illustrated by different colours. The *y-axis* indicates the Bayes Factor corrected by local score method (Lindley score).

Interestingly, the genomic architecture associated with *B. incana* — pollinator interaction indices was slightly more complex, with the detection of multiple narrow peaks per variable (**Fig. 3, Fig. S4**). As expected from their ecological correlations, some indices describing *B. incana* - pollinator interactions shared genomic regions among them (**Fig. 4 and Fig. S5**). Considering all the indices related to *B. incana* – pollinator interactions, 73% of SNPs were associated with the response of *B. incana* to the variation of these indices (**Fig. S6**). The remaining SNPs were shared among these indices and the variables involved in adaptive response to mean annual temperature, texture of the soil (fine silt and coarse sand) and some functional categories of pollinators (**Fig. S5**). Finally, 92% and 85% of SNPs with highest association scores were uniquely associated with the adaptation of *B. incana* to soil and climate respectively (**Fig. S6**). Overall, our results highlighted a flexible genetic architecture involving mainly unique genomic regions in the adaptive response to ecological variables, as well as few shared genomic regions among them.

**Figure 4.**
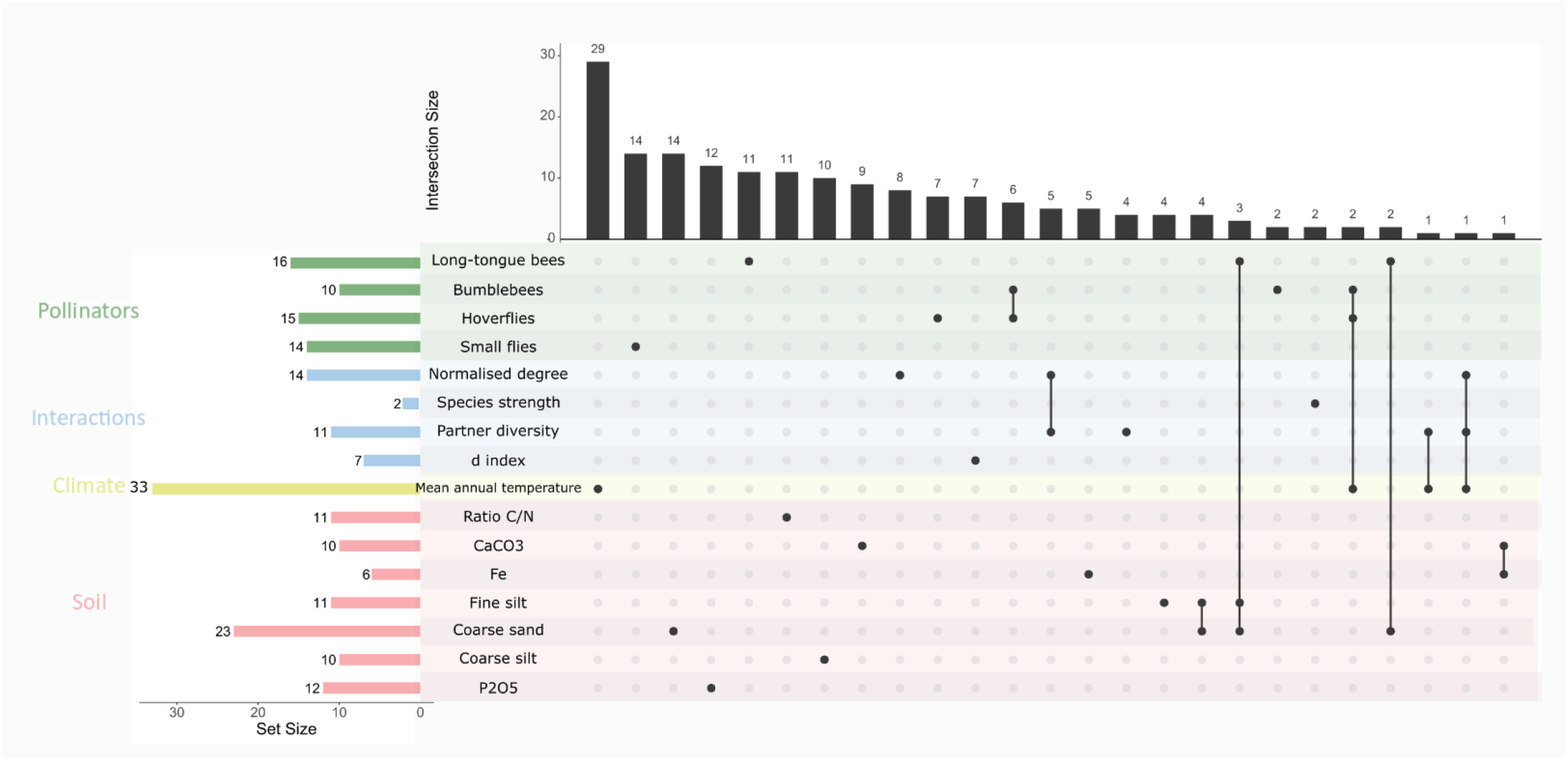
Illustration of the relationship among candidate genes associated with local adaptation to ecological network. Only variables for which a significant enrichment of the selection signature was detected are considered. The left shows the number of candidate genes (set size) identified in local adaptation to the specific variable in GEA analysis. On top, the number of candidate genes associated with a specific variable (single black dot) or shared among variables (multiple black dots linked). The candidate genes are those from the significant zones identified by correcting with local score method the GEA, and the down and upstream genes.

### Local adaptation of *Brassica incana* to ecological network

To address the signal of natural selection on loci identified by GEA, we performed a genome-wide scan for genetic differentiation index (XtX) among the 21 natural populations of *B. incana* based on standardized allelic frequencies (*i*.*e*. allele frequencies corrected for population structure). After correcting the signal with local score method, we detected four genomic regions under strong selection on super-scaffolds 1 (including five candidate genes), 10 (including three candidate genes), 37 (including nine candidate genes) and 74 (including 32 candidate genes, **Fig. S7, Table S8**).

To support local adaptation of plants to different ecological variables, we performed an enrichment for signature of selection by testing the over-representation of top SNPs (i.e. 0.05% upper tail of BFdB distribution of SNPs associated with the 33 ecological variables) in the tail of the XtX distribution (i.e. 0.05% of SNPs strongly under selection). The enrichment in signature of selection allowed to distinguish the ecological variables associated with genomic variations due to selective processes from those that were associated due to random effects. We found that 17 out of 33 ecological variables displayed a significant enrichment (**Table 1, Table S7**). For instance, we found a strong enrichment in signature of selection for five functional categories of pollinators including bumblebees (23-fold), hoverflies (25-fold), and long-tongue bees (21-fold, **Table 1**). The four variables involved in *B. incana* — pollinator interactions showed significant enrichment ranging from 6-fold for species strength to 31 for the d index (**Table 1**). Finally, abiotic factors showed significant fold-enrichment for 7 out of 15 edaphic variables (ranging from 12 for CaCO3 to 62 for coarse sand), and a strong significant enrichment for mean annual temperature (109-fold, **Table 1, Table S7**).

**Table 1.** Significant enrichment in signature of selection for pollinator categories, plant-pollinators interaction indices, climatic and edaphic variables testing the over-representation of the 0.05% upper tail of the Lindley score distribution in the 0.05% upper tail of the genome-wide spatial differentiation (XtX) distribution. See Table S7 for the enrichment’s results of the 33 ecological variables.

| Traits | ntops | Enrichment | pvalue |
| --- | --- | --- | --- |
| Long-tongue bees | 21 | 16.53 | ** |
| Bumblebees | 23 | 18.11 | *** |
| Large bees | 15 | 11.81 | ** |
| Hoverflies | 25 | 19.68 | *** |
| Small flies | 6 | 4.72 | * |
| Normalised degree | 19 | 14.96 | ** |
| Species strength | 6 | 4.72 | * |
| Partner diversity | 20 | 15.74 | *** |
| d index | 31 | 24.40 | *** |
| Mean annual temperature | 109 | 85.81 | *** |
| Ratio C/N | 13 | 10.23 | ** |
| CaCO3 | 12 | 9.45 | ** |
| Fe | 39 | 30.70 | *** |
| Fine silt | 59 | 46.45 | *** |
| Coarse sand | 62 | 48.81 | ** |
| Coarse silt | 31 | 24.40 | ** |
| P2O5 | 15 | 11.81 | ** |

### Candidate genes involved in plant response to pollinators

The candidate genes involved in local adaptation of *B. incana* to the ecological network were identified by retrieving genes within significant zones identified by the GEA analyses, as well as down- and upstream genes as in Libourel et al. (2021). From the GEA results showing significant enrichment in XtX index, we identified 48 candidate genes involved in plant responses to functional categories of pollinators, and 26 candidate genes involved in adaptive responses to *B. incana* — pollinator interactions (list of all candidate genes in **Table S8**). For candidate genes involved in adaptation to different pollinator functional categories, we found some genes involved in plant signals and rewards such as (1) UV-B-induced protein inducing changes in the accumulation of phenolic compounds, carotenoids and glucosinolates (Schreiner et al. 2012), (2) Sinapine esterase, an important enzyme in floral pigmentation (Nguyen et al. 2021), (3) 2-C-methyl-D-erythritol 2,4-cyclodiphosphate synthase (*ISPF*) involved in methylerythritol phosphate pathway (MEP) in biosynthesis of terpenoids (Tarkowska- and Strnad 2018), an important volatile class involved in plant attractiveness (Abas et 2017, Bouwmeester et al 2019), (4) Dihydropyrimidine dehydrogenase (NADP(+)) (*PYD1*) involved in β-alanine biosynthesis (Wang et al 2021) a component present in nectar-feeding bumblebees (Rossi et al. 2014), (5) Trehalose-phosphate phosphatase B (*TPPB*) important in carbon flux maintain associated with sucrose supply (Nunes et al. 2013), a main component of nectar, and (6) transcription factor MYB73 involved in anthocyanin biosynthetic pathway (Gomez et al 2020). We found several candidate genes involved in plant architecture and growth such as (1) protein CUP-SHAPED COTYLEDON 3 (*NAC031*) (Gao et al. 2021), and (2) nuclear pore complex protein (*NUP98A*) (Parry 2014), (3) Pectinesterase inhibitor 10 (*PMEI10*) (Wormit et al 2018). Some candidate genes were involved in reproduction processes such as (1) Expansin-B5 in growth pollen tube (*EXPB5*, da Costa et al 2012), and (2) type I inositol polyphosphate 5-phosphatase 12 (*IP5P12*) involved in pollen dormancy and early germination (Wang et al 2012). We identified candidate genes involved in immunity and plant defence such as Serine/threonine-protein kinase *BSK7* interacting with pattern-triggered immunity (Majhi et al 2021), and LRR receptor-like serine/threonine-protein kinase (Afzal et al 2008). It is noteworthy that 38% of the identified candidate genes involved in plant response to pollinators were associated with proteins with unknown function. Interesting, few candidate genes mentioned above were involved in both the response of *B. incana* to bumblebees and hoverflies such as UV-B induced protein, or the PYD1 (**Fig. 4, Table S8**).

For candidate genes involved in adaptation to plant-pollinator interactions, we found some genes involved in plant architecture and growth such as transcription factor BEE2 (*BEE2*) (Friedrichsen et al 2002), protein TRIGALACTOSYLDIACYLGLYCEROL 2 (TGD2)(Fan et al. 2015), and ethylene-responsive transcription factor (*ERF024*, Lata et al 2014). Some of these candidate genes were involved in controlling pollen tube such as *LLG3* GPI-anchored protein *LLG2* (Ge et al. 2019), or in flowering time such as polyadenylation and cleavage factor homolog 4 (*PCFS4*) (Xing et al 2008). However, 48% of the identified candidate genes involved in *B. incana* response to plant-pollinator interactions are associated with proteins with unknown functions.

Finally, we observed multiple shared candidate genes in the response of *B. incana* to pollinators and climatic factors: between mean annual temperature and hoverfly and bumblebee visitation, pollinator diversity, and combined pollinator diversity and realized number of pollinator links, as well as shared candidate genes in the response of *B. incana* to pollinators and edaphic factors, *e*.*g*. between long-tongue bees and coarse sand, and coarse sand and fine silt (**Fig. 4, Table S8**).

## Discussion

While pollinators provide essential ecosystem services (Klein et al. 2007, Potts et al. 2010), whether and how plants with generalized pollination adapt to geographic variation in pollinator communities and the underlying genetic basis is still poorly documented. Using an ecological genomics approach, our study unravelled the genomic bases of plant adaptation to pollinator communities and potential interacting abiotic factors.

### Local adaptation to functional categories of pollinators

The observed mosaic of pollinators among our 21 natural populations led to local adaptation of the generalist plant species *B. incana* to pollinators. These results are in line with few studies in evolutionary ecology emphasizing the importance of pollinators in driving the floral evolution of generalist plant species (Gomez et al. 2009, Bodbyl Roels and Kelly 2011, Sahli and Conner 2011, Gomez et al. 2015, Sobral et al. 2015, Gervasi and Schiestl 2017, Schiestl et al. 2018, de Manincor et al. 2021). We uncovered here the underlying genomic mechanisms of these adaptive processes: a flexible genomic architecture involving genomic regions that were strongly associated with functional categories of pollinators. In particular, we have identified pollinator category-specific candidate genes including some that were potentially involved in biosynthetic pathways of plant signals and rewards to attract pollinators. For instance, we identified two interesting genes involved in *B. incana* adaptive response to long-tongue bees; a candidate gene encoding for the enzyme 2-C-methyl-D-erythritol 2,4-cyclodiphosphate synthase (*ISPF*) involved in the ethylerythritol phosphate (*MEP*) pathway, responsible for terpenoids biosynthesis (mono- and diterpenoids biosynthesis; Abbas et al. 2017, Tarkowská and Strnad 2018, Bouwmeester et al. 2019), an important class of volatiles in plant-pollinator interactions (Baldwin et al. 2006, Abas et al. 2017, Bouwmeester et al. 2019). In addition, we found a candidate gene encoding for the trehalose-phosphate phosphatase B (*TPPB*) enzyme involved in carbon flux maintaining, correlated with sucrose supply (Nunes et al. 2013), an essential component of nectar. Interestingly, we found adaptive responses of *B. incana* both to efficient pollinators in pollen transfer (long-tongue bees, bumblebees, and other large bees), as well as to supposedly less efficient pollinators (hoverflies, small flies). Local adaptation to supposedly “inefficient” pollinators may be surprising since they contribute less to plant reproductive success. However, the observed adaptation to such less-efficient pollinators can be explained by spatial and temporal variation in selective regimes influenced by the local interactions (Gomez et al. 2009b) such as increased importance of hoverflies to pollination when bees are scarce or absent (Jauker and Wolters 2008, Ohashi et al. 2021). Local adaptation to less-efficient pollinators could also be related to reproductive assurance in hoverfly-dominated populations, with potential limited pollen transfer leading to reduced floral size and decrease of volatile emission as observed in Gervasi and Schiestl (2017) or related to indirect effects of hoverflies on bees’ visitations. Surprisingly, in our study, it appears that the adaptive response to bumblebees and hoverflies involved similar genomic regions. For instance, we identified a candidate gene encoding for a dihydropyrimidine dehydrogenase (*PYD1*) enzyme involved in the biosynthesis of β-alanine (Wang et al 2021), a component recently found associated with nectar-feeding bumblebees in *Gentiana lutea* (Rossi et al. 2014). It would be interesting to compare the rate of β-alanine production among our populations according to the ratio of bumblebees and hoverflies visitations to better understand this adaptive response. Finally, among the identified candidate genes involved in response to different functional categories of pollinators, we also identified genes involved in protein chaperon, plant growth, plant immunity, and a non negligeable part of protein with unknown function (∼33% of candidate genes). Thus, plant species with generalized pollinations system are locally adapted to their “specific” generalist pollinator community including identifiable genomic regions associated with candidate genes that are involved in plant interactions with specific pollinator functional categories.

### Local adaptation to ecological networks

We demonstrated that *B. incana* is able to adapt simultaneously to different stimuli driving local adaptation to complex ecological networks. First, by interacting with a broad and diverse community of pollinators, the adaptative responses of generalist species have long been considered unlikely due to simultaneous conflicting and highly variable selective pressures imposed on flowers. However, we demonstrated genome-wide signatures of adaptation for multispecies assemblage of pollinators. This agrees with previous studies in evolutionary ecology showing the adaptive response of generalist species to pollinator communities (Gomez et al 2009a, Gomez et al 2009b, Sahli and Conner 2011, Lomascola et al. 2019). Our results also highlighted a lack of knowledge on molecular mechanisms involved in this adaptive process because 48% of the identified candidate genes were associated with proteins of unknown function. Interestingly, the genomic architecture underlying the response of *B. incana* to the assemblage of pollinators was not the sum of genetic variants specific to functional categories of pollinators indicating nonadditive selection acting on *B. incana* as previously observed in plant-plant interactions (Baron et al. 2015, Libourel et al. 2021). This assumption agrees with a previous evolutionary study in *B. rapa* observing nonadditive selection for floral traits where phenotypic evolution mediated by the combination of two pollinator species was different from that mediated by either pollinator in isolation (Schiestl et al. 2018). A phenotypic characterisation of our populations is still needed to better understand this evolutionary process. Nonadditive selection seems to be a common process in natural populations caused by indirect ecological effects. Such effects remain unpredictable by pairwise selection, and difficult to study due to infinite number of ecological factors to be considered (Sahli and Conner 2011, Terhorst et al. 2015). To estimate potential indirect effects from abiotic factors on plant-pollinator interactions, we compared shared genetic variants among adaptive responses to abiotic and biotic factors. We observed only few shared candidate genes involved in the local adaptation of *B. incana* to long-tongue bees, structure of the soil and temperature. Long-tongued bees, including the genus *Anthophora* in our study, are ground-nesting bees, hence variation in soil texture could have a significant impact on their occurrence in populations (Antoine and Forrest 2021). However, further ecological characterisation is needed to control indirect effects such as local climate, composition of lower soil layers, microbiome, herbivores, or surrounding flowering plants for instance. In addition, due to the geology of Southern Italy, we had a strong confounding effect between population structure (controlled by Bayesian model) and the type of soil (following a Northwest – Southeast axis) likely leading to an underestimation of adaptive potential to soil with the presence of false negatives in our GEA analysis and underestimating the role of soil in plant-pollinator interactions. By illustrating the effect of complex ecological network on generalist plants through a flexible genomic architecture, our results highlighted the importance of considering ecological variables in the adaptative landscape of generalist species and abiotic factors to better understand their impact on plant evolution (Carvalheiro et al 2021). With the current declines in insect diversity and its potential impact on flowering plant reproductive success, we stress the need to expand our knowledge of the adaptive potential of plants to pollinator communities using a multi-disciplinary approach from ecology to molecular biology to genomics.

## Methods

### Natural populations of *Brassica incana*

We used the non-model plant species *Brassica incana* (**Fig. 1**), an allogamous and self-incompatible perennial species growing on cliffs, and mainly distributed in Southern Italy (Landucci et al. 2014, Ciancaleoni *et al*. 2018). This wild species is a close relative species of *Brassica oleracea* crop species (Landucci et al. 2014, El-Esawi 2017). From the data available in the literature and our own observations, we found 40 populations in Southern Italy. We used 21 natural populations (**fig. 1, table S1**) for which at least 20 individuals were present in spring 2018 with safe access. Our populations grew on two distinct types of soil: six populations on tuff soil and 15 populations on limestone soil (**fig. 1, table S1**). The populations were located from 2m to 767m elevation (average = 278m, **Table S1**), with an average distance of 61.78 km (median = 41.9 km, minimum = 1.25 km, maximum = 168.6 km).

### Ecological characterization

We characterized the soil of the 21 natural populations of *B. incana* during spring 2018 by collecting two samples per population on the ground surface (maximum depth ∼10cm). The samples were sent to the Soil Analysis Laboratory of Arras (INRA, France, https://www6.hautsdefrance.inrae.fr/las). Twenty-one soil compounds were measured (**Dataset1**): aluminium (Al), carbon (C), ration carbon/nitrogen (ratio C/N), calcium (Ca), total calcium carbonate (CaCO3), clay (< 0 µm), total copper (Cu), iron (Fe), fine sand (0.05mm to 0.2mm), coarse sand (0.2mm to 2mm), fine silt (2µm to 20 µm), coarse silt (20µm to 50 µm), potassium (K), magnesium (Mg), manganese (Mn), total nitrogen (N), sodium (Na), organic matter (om), phosphorus (P2O5), silicon (Si) and zinc (Zn). We followed the same method as described in Brachi et al. 2013, and all protocols are available at https://www6.hautsdefrance.inrae.fr/las/Methodes-d-analyse/Sols.

We retrieved 20 biologically meaningful climatic variables (**Dataset1**) for the 21 populations from ClimateEU database (v4.63 software, described in Hamman et al. 2013). Like Frachon et al. (2018), the average data across 2003-2013 were used for these 21 climatic data related to temperature and precipitation.

We characterized pollinator communities in spring 2018 (for 17 out of 21 populations) and spring 2019 (for 19 out 21 populations) for a total of 19 biotic variables (**Dataset1**). To fully characterize the pollinator communities, one to four sessions (in average three sessions) of observations were conducted in spring 2018 and 2019 (one session in 2018, and three in 2019). We recorded pollinators’ visits for one hour starting at 11.30 am in each population, with approximately observations of 10 minutes per plant within the population. In average, five plants were observed per session (median = 5 plants, maximum = 11 plants, minimum = 1 plant). We assigned visitors into 13 functional categories: bumblebees, long-tongue bees (genus Anthophora), other large bees (called large bees), small bees, honeybees, large wasps, small flies, large flies, hoverflies, small beetles, large beetles, butterflies, and wasps. Due to the scares number of wasp visits (only one visit in CHIU population), it was discarded from the dataset, except for the plant-flower visitor network.

Because our study was population-centred, we estimated Best Linear Unbiased Predictions (BLUP) *i*.*e*. the average pollinator visits per population using a mixed model in the R Studio environment (package lme4, Bates et al. 2015).

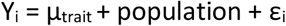

where Y_i_ is BLUP for visits by functional categories of pollinators, µ_trait_ the overall average of the trait (observed number of visits of functional categories of pollinators), population is considered as random effect, and ε_i_ is the residual variance.

The plant-flower visitor network was constructed using the *bipartite* package (Dormann *et al*., 2008) based on the total number of visits within the populations from the 12 distinct functional categories of pollinators among the 21 natural populations of *B. incana*. Similarly to the species-species interaction networks, category-level indices for each population were calculated using *bipartite* (Dormann, 2011). We calculated eight following indices as described in Dormann *et al*. 2011 and called latter *B. incana* — pollinator interaction metrics: (1) normalised degree representing the number of partner species in relation to the potential number of partner species, (2) species strength representing the sum of dependencies of each species, aiming at quantifying a species’ relevance across all its partners, (3) species specificity index representing the coefficient of variation of interactions, normalised to values between 0 (low variability suggesting low specificity) and 1 (high variability suggesting high specificity), (4) partner diversity representing the Shannon diversity index of the interactions of each species, (5) effective partners representing the logbase to the power of “partner diversity” interpreting as the effective number of partners, if each partner was equally common, (6) proportional similarity representing the specialization measured as dissimilarity between resource use and availability, (7) proportional generality representing the effective partners’ divided by effective number of resources; this is the quantitative version of proportional resource use or normalised degree (i.e., the number of partner species in relation to the potential number of partner species), and (8) d index representing the specialisation of each species based on its discrimination from random selection of partners.

The matrix of spearman correlation for the 61 ecological variables (11 functional categories of pollinators, eight *B. incana* interaction indices, 20 climatic variables, and 22 edaphic variables) using the R package Hmisc (Harrell et al. 2021). We pruned the set of variables using the pairwise Spearman correlations among variables, and only variables with spearman’s rho < 0.8 were retained for the genomic analysis. In total, we kept 33 ecological variables: 11 functional categories of pollinators, four *B. incana* — pollinator interaction indices, three climatic variables, and 15 edaphic variables. We performed a principal component analysis representing the distribution of 33 ecological variables among the 21 populations using ade4 package in R (Dray and Dufour 2007).

### *De novo* reference genome

#### DNA extraction

We chose one individual from the island of Capri population as reference genome (CAPR in **fig. 1**), a stable population between 1984 and 2012 with low gene flow with cultivated plants (Ciancaleoni et al. 2018). Seeds from Capri population were collected in 2017, sown in a phytotron in the summer of 2018 (24 hours light, 21°C, 60% humidity, watered twice a day), and grown in air-conditioned greenhouse at the University of Zürich in standard condition (22.5°C, 50-60% of humidity, additional light). Prior to DNA extraction, plants were kept in the dark for two days, which reduced the amount of polysaccharides that interfere with the DNA extraction yield. We modified the high-molecular weight genomic DNA extraction protocol from Mayjonade *et al*. (2016) as described in Russo et al. (2022). Briefly, this extraction was performed in 23 parallel tubes to increase the quantity of final DNA. The 23 DNA extracts were pooled together, and purifying using carboxylated magnetic beads as explained in Mayjonade et al. (2016) protocol. We measured 146ng/µL of total DNA concentration using a nanodrop (ratio A-260/A-280 = 1.85, ratio A-260/A-230 = 2.19) and 152ng/ µL with Qubit. The purified sample was sent to the Functional Genomic Center of Zürich (FGCZ) for library preparation and three different next generation sequencing were performed to obtain a *de novo* reference genome of *Brassica incana*.

#### PacBio library preparation and sequencing

The continuous long read (CLR) SMRT bell library was produced using the SMRTbell Express Template Prep Kit 1.0. (Pacific Biosciences) at the functional genomic center Zürich (FGCZ). The input genomic DNA concentration was measured using a Qubit Fluorometer dsDNA Broad Range assay (Thermo). The high molecular weight (HMW) genomic DNA (gDNA) sample (6 μg) was mechanically sheared to an average size distribution of 30 kbp, using a g-TUBE (Covaris) on a minispin plus centrifuge (Eppendorf). A Femto Pulse gDNA analysis assay (Agilent) was used to assess the fragment size distribution. Sheared gDNA was DNA damage repaired and end-repaired using polishing enzymes. PacBio sequencing adapters were ligated to the DNA template, according to the manufacturer’s instructions. A Blue Pippin device (Sage Science) was used to size select the SMRT bell library and enrich for fragments > 25 kbp. The size selected library was quality inspected and quantified using a Femto Pulse gDNA analysis assay (Agilent) and on a Qubit Fluorimeter (Thermo) respectively. A ready to sequence SMRT bell-Polymerase Complex was created using the Sequel binding kit 3.0 (Pacific Biosciences P/N 101- 500-400) according to the manufacturer instructions. The Pacific Biosciences Sequel instrument was programmed to sequence the library on five Sequel™ SMRT® Cells 1M v3 (Pacific Biosciences), taking one movie of 10 hours per cell, using the Sequel Sequencing Kit 3.0 (Pacific Biosciences). After the run, the sequencing data quality was checked, via the PacBio SMRT Link software (v 6.0.0.47841), using the “run QC module” (**Table S2**).

#### Illumina library preparation and sequencing

The TruSeq DNA Nano Sample Prep Kit v2 (Illumina, Inc, California, USA) was used in the succeeding steps. DNA samples (100 ng) were sonicated with the Covaris using settings specific to the fragment size of 350 bp. The fragmented DNA samples were size-selected using AMpure beads, end-repaired and adenylated. TruSeq adapters containing Unique Dual Indices (UDI) for multiplexing were ligated to the size-selected DNA samples. Fragments containing TruSeq adapters on both ends were selectively enriched by Polymerase chain reaction (PCR). The quality and quantity of the enriched libraries were validated using Tapestation (Agilent, Waldbronn, Germany). The product was a smear with an average fragment size of approximately 500 bp. The libraries were normalized to 10nM in Tris-Cl 10 mM, pH8.5 with 0.1% Tween 20. The Novaseq 6000 (Illumina, Inc, California, USA) was used for cluster generation and sequencing according to standard protocol. Sequencing was paired end (PE) at 2 X150 bp. This described protocol was used for both *de novo* sequencing of the reference individual, as well as the Pool-sequencing of the 21 natural populations.

#### Pre-processing and Mapping of Illumina Reads

Quality control and Bowtie2 alignment of the Illumina PE reads were performed using data analysis workflows in the R-meta package ezRun (https://github.com/uzh/ezRun), managed by the data analysis framework SUSHI (Hatakeyama *et al*., 2016), which was developed and maintained by FGCZ. Technical quality was evaluated using FastQC (v0.11.7). We screened for possible contaminations using FastqScreen (v0.11.1) against a customized database in ezRun, which consists of SILVA rRNA sequences (https://www.arb-silva.de/), UniVec (https://www.ncbi.nlm.nih.gov/tools/vecscreen/univec/) sequences, refseq mRNA sequences and selected refseq genome sequences (human, mouse, *Arabidopsis*, bacteria, virus, phix, lambda, and mycoplasma) (https://www.ncbi.nlm.nih.gov/refseq/). Illumina PE reads were pre-processed using fastp (v0.20.0), with which sequencing adapters and low-quality ends (4 bp sliding windows from both ends, average quality < Q20) were trimmed. Trimmed reads passing the filtering criteria (average quality >= Q20, minimum length >=18 bp) were aligned using Bowtie2 (v2.4.1) with the “--very-sensitive” option. Trimmed reads from the reference individual were aligned to the PacBio HG4P4 assembled contigs for genome polishing. Afterwards trimmed reads from the 21 natural populations were aligned to the polished and scaffolded genome assembly for variant analysis. PCR-duplicates were marked using Picard (v2.18.0). Read alignments were comprehensively evaluated using the mapping QC app in ezRun in terms of different aspects of DNA-seq experiments, such as sequence and mapping quality, sequencing depth, coverage uniformity and read distribution over the genome (**Table S2**).

#### De novo Genome Assembly

PacBio subreads from all five SMRT cells were merged and assembled using HGAP4 (Hierarchical Genome Assembly Process v4) in the PacBio SMRT Link software (v 6.0.0.47841). Before being assembled, subreads were filtered with read quality of 70%. The estimated genome size was set at 650 Mbp. Illumina PE reads from the same sample were pre-processed and mapped to the assembled primary contigs as described above. Assembled primary contig sequences were then further polished with mapped Illumina PE reads using pilon (v1.23). Only reads with mapping quality above Q20 and bases with phred scores above Q20 were used for the polishing.

#### In silico genome digestion and Bionano Optical Mapping

The polished genome assembly was first *in silico* digested using Bionano Access software (v1.2.1) to evaluate whether the nicking enzyme (Nb.BspQI), with recognition sequence GCTCTTC, and the non-nicking enzyme DLE-1, with recognition sequence CTTAAG, were suitable for optical mapping in the genome. An average of 13.6 nicks/100 kbp with a nick-to-nick distance N50 of 13,734 bp was expected for Nb.BspQI, while DLE-1 was found to induce 22.2 nicks/100 kbp with a nick-to-nick distance N50 of 8,054 bp. The values were in line with manufacturer’s requirements.

For the Direct Label and Stain (DLS) protocol, the DNA sample was labelled using the Bionano Prep DNA Labeling Kit-DLS (cat. no. 80005) according to manufacturer’s instructions. In details, 750 ng of purified gDNA was labelled with DLE-1 labelling mix and subsequently incubated with Proteinase K (Qiagen, cat. no. 158920) followed by drop dialysis. After the clean-up step, the DNA was pre-stained, homogenized, and quantified using on a Qubit Fluorometer to establish the appropriate amount of backbone stain. The reaction was incubated at room temperature for at least 2 hours. For the Nick Label Repair and Stain (NLRS) protocol, the DNA sample was labelled using the Bionano Prep DNA Labelling Kit-NLRS according to manufacturer’s instructions (Bionano Genomics, cat. no. 80001). In details, 300 ng of purified gDNA was nicked with Nb.BspQI (New England BioLabs, cat. no. R0644S) in NEB Buffer 3. The nicked DNA was labelled with a fluorescent-dUTP nucleotide analogue using Taq DNA polymerase (New England BioLabs, cat. no. M0267S). After labelling, nicks were ligated with Taq DNA ligase (New England BioLabs, cat. no. M0208S) in the presence of dNTPs. The backbone of fluorescently labelled DNA was counterstained overnight with YOYO-1 (Bionano Genomics, cat. no. 80001). DLS and NLRS labelled DNA samples were loaded into a nanochannel array of a Saphyr Chip (Bionano Genomics, cat. no. FC-030-01) and run by electrophoresis each into a compartment. Linearized DNA molecules were imaged using the Saphyr system and associated software (Bionano Genomics, cat. no. 90001 and CR-002-01). BioNano row molecule data are available on **Table S3**.

#### Assembly of Optical Maps and Hybrid Scaffolding

The *de novo* assembly of the optical maps was performed using the Bionano Access (v1.2.1) and Bionano Solve (v3.2.1) software. The assembly type performed was the “Saphyr data”, “non-human”, “non-haplotype” with “extend and split” and “cut segdups”. Default parameters were adjusted to accommodate the genomic properties of the *Brassica incana* genome. Specifically, the “Initial P value” cut-off threshold was adjusted to 1 × 10^−10^ and the P value cut-off threshold for extension and refinement was set to 1 × 10^−11^ according to manufacturer’s guidelines (default values are 1 × 10^−11^ and 1 × 10^−12^, respectively). Dual-enzyme hybrid scaffolding was then performed using the same software suits with default parameters. This dual-enzyme hybrid scaffolding used the Bionano optical maps to scaffold polished (PacBio and Illumina) contigs.

#### Genome Annotation

Repeat sequences in the *de novo* assembled genome were predicted using RepeatScount (v1.0.6). Predicted repeat sequences and known transposable elements (TEs) in *Brassica oleracea* were masked using RepeatMasker (v4.1). Gene model prediction was performed using maker (v3.01.03). In details, *ab initio* gene prediction was performed using AUGUSTUS with the pre-trained parameter set for *Arabidopsis*. Protein and cDNA sequences of *B. oleracea* (Ensemble release 42) were aligned to the assembled genome and used as supporting evidence for gene prediction. For functional annotation, prediction protein sequences were compared to the SwissProt database (release 2019_03) using blastp (v2.6.0), and the InterPro database using interproscan (v5.32-71.0).

### Genomic characterization of 21 populations using a pool-sequencing approach

In spring 2018, we collected leave tissue from, in average, 28 individuals per population (median = 30 plants, max = 30 plants, min = 15 plants, *i*.*e*. a total of 590 samples) in 1.5mL Eppendorf tubes. The samples were stored during the field day in dry ice and moved into -80°C freezer at the end of field day. The DNA extraction was performed in fall 2018 by grinding samples using two beads, cooling down in liquid nitrogen, and crushed them with 30 vibrations/second three times 30 seconds. We extracted DNA using the sbeadex® maxi plant kit from LGC Genomics in Kingfisher™ Flex Purification Systems (Thermo Scientific™), a magnetic-particle robot at the Genetic Diversity Centre (GDC) Zürich platform. We added 250µL of lysis buffer in all homogenised samples. After homogenization (2-3 seconds on vortex, and 20 reversing tubes), we incubated our samples 20 minutes at 65°C. We added 1.12µL of RNAse (940U/mL) and reversing tubes 10 times. We incubated the samples 10 more minutes at 65°C. After centrifuging at 2.5×1000 rcf for 10 minutes at 20°C, we transferred 200µL of the lysate in deep 96-well plates with 520µL of binding buffer and 60µL of sbeadex particles suspension. The samples were incorporated into the Kingfisher™ robot for the DNA purification. After bringing magnets into contact with the tubes for 1 minutes, the supernatant was removed and discarded. 400µL of wash buffer PN1 was added in each sample and mixed by pipetting to re-suspend the pellet. After 10 minutes of incubation and agitation at room temperature, the magnets were brought into contact with the tubes for 1 minute. The supernatant was removed and discarded, and a second round of washing was performed adding 400µL of wash buffer PN2 in each sample, incubating 10 minutes at room temperature and bringing the magnets into contact with tubes. The supernatant was removed and discarded. 100µL of elution buffer PN was added to the pellet and mixed by pipetting. The solution incubated at 55°C for 10 minutes, and finally the magnets were brought into contact with tubes for 3 minutes until the sbeadex formed a pellet and stayed on the magnets. The eluate of the samples was transfer to a new 96-well plates and stored in the fridge.

The DNA concentration of all samples were measured using ddDNA Qubit assay measurement on plate reader Spark M10 (excitation wavelength = 485nm, emission wavelength = 535nm). Eight samples with too low DNA concentration were discarded. In total 582 samples were used for the pool sequencing with a DNA concentration superior to 1.5 ng/µL (average = 12.68 ng/µL, median = 10.18 ng/µL, max = 61.09 ng/µL). For each of the 21 populations, the individuals were pooled together equimolarly, with an average of 27.7 individual per pool (median = 29 individuals, minimum = 15 individuals, maximum = 30 individuals).

We proceeded for the pool-sequencing as previously described in the methods for the *de novo* reference genome sequencing using Illumina sequencing.

#### Freebayes variant calling

Multi-samples frequency-based (-F 0.05) variant calls (--use-best-n-alleles 4 --pooled-continuous) were generated using the freebayes-parallel script in freebayes (v1.2.0-4-gd15209e, Garrison and Marth 2012), with 16 threads of freebayes running in parallel across regions of 100kb in the *de novo* polished genome assembly (PacBio, Illumina and Bionano). Single nucleotide polymorphisms (SNPs) with variant quality above Q20 were retained using bcftool (v1.9) for downstream analysis and were annotated with *de novo* predicted gene models using SnpEff (v4.2). The final dataset was composed of 6’899’774 SNPs across the 21 natural populations of *B. incana*.

### Data filtering

The matrix of population allele frequencies was trimmed using VCFtools (Danecek et al. 2011) and following Frachon et al. (2018). We kept only biallelic loci (391’671 SNPs discarded) and removed the indels (7’960 SNPs discarded). We discarded SNPs with a minimum mean read depth lower than 6, and higher than 100 (143’710 SNPs discarded). We removed all SNPs with missing value in more than two populations (613’387 SNPs discarded). We finally kept only 139 super-scaffolds (203’955 SNPs discarded). The final allele read count matrix included 5’530’708 SNPs for 21 populations.

### Genome Environment Association (GEA) analysis on 33 ecological variables

We performed a GEA analysis using a pool-sequencing approach between the 5’530’708 SNPs and 15 variables describing pollinator communities (11 functional categories of pollinators, and 4 *B. incana* – pollinators’ interactions), 3 climatic variables and 15 edaphic variables. Genome scans were based on Bayesian hierarchical model implemented in Baypass software (Gautier 2015). Considering the covariance matrix of allele frequencies among population, this model allowed to correct potential effect of demographic histories (Gautier 2015). As described in Frachon et al. (2019), we used the core model to estimate the Bayesian Factor (BF_is_ in dB) between the allelic frequencies along the genome, and different descriptors of pollinator communities as well as abiotic variables. The core model was repeated three times due to Importance Sampling algorithm, and the final Bayesian Factor was estimated by averaging them. Considering the large amount of SNPs used, we sub-sampled the procedure to estimate the matrix of population allele frequencies (Ω) as in Frachon et al. (2018), by dividing the full data set into 19 sub-data sets of ∼ 254’785 SNPs each. The GEA for each trait and each sub-data set were performed in parallel and merged again after analyses. Finally, we corrected the Bayes factor (BF_is_ in dB called later BFdB) obtained by using a local score approach to consider the linkage disequilibrium (Bonhomme et al. 2019) allowing to detect the accumulation of similar *p-value* in the same region increasing the power of genomic analyses. To do, we artificially created *p-values* by ranking the BFdB value from the highest to the smallest ones and divided the rank by the total number of SNPs. The parameter ξ was fixed at three for the local score method (Bonhomme et al. 2019, Libourel et al. 2021). We used upset plots to detect shared SNPs and candidate genes among the 33 ecological variables considering 0.05% SNPs with highest association score after local score method (R package UpSetR, Gehlenborg et al. 2019). Due to the geology of Southern Italy, we observed a Northwest-Southeast axis of variation of type of soil (tuff *versus* limestone), potentially matching the demographic history. Because GEA analyses are based on corrected allele frequency by population structure, we may observe false negative. We estimated the genomic variation among the population using a singular value decomposition (SVD) of the matrix of raw allele frequency (without population structure correction). A significant correlation between the genomic variation from SVD and environmental variable would indicate the presence of potential false negative.

### Signature of selection

We performed a genome-wide selection scan among the 21 populations based on the XtX spatial genetic differentiation (Günther and Coop 2013, Gautier 2015). This index considered the standardized allele frequencies of a given SNP, a measure of the variance of allele frequencies across 21 natural populations. This method has been demonstrated to be successful for natural populations (Frachon et al. 2018, Frachon et al. 2019). As described above, we also implemented the local score approach to correct the XtX fixing parameter ξ at three. Finally, we estimated the enrichment in signature of selection by testing whether the SNPs with the highest association scores with environmental variables (0.05% upper tail of the local score) were significantly enriched in the 0.05% upper tail of XtX distribution (Brachi et al. 2015, Frachon et al. 2018, Frachon et al. 2019). The significance of the enrichment was testing using the method described in Hancock et al. (2011) by running 10’000 null circular permutations of the 0.05% SNPs with highest association score with 33 environmental variables.

### Identification of candidate genes

To identity candidate genes involved in local adaptation of *B. incana* to pollinator communities and abiotic variables, we retrieved genes within the significant zone identified by the GEA analyses and corrected by the local score approach, and down and upstream genes of these zones as in Libourel et al. (2021). Only zones containing more than 3 SNPs were kept.

## Supporting information

Table S8 candidate genes

## Acknowledgement

We are grateful to David Preiswerk, Cesario Capasso and Samson Accoca-Pidolle for assisting us with the field experiment. We thank Cyril Libourel for discussion regarding genome annotation and improvement of gene identification. We thank Anne Roulin for her comments on a previous version of the draft. DNA extraction and pooling samples in this paper were performed in collaboration with the Genetic Diversity Centre (GDC), ETH Zurich. We thank the Functional Genomic Centre of Zürich (FGCZ) for their support in library preparation and sequencing. This research was funded by the Swiss National Science Funds (SNF grant no. 31003A_172988 to F.P.S.). In addition, University of Naples (UniNA) in the framework of Program STAR-GENPOLL, and the University of Zürich provided funding.

## Data availability

All data will be available after acceptance of the manuscript as described hereafter. Sequencing data from Pacbio and Illumina used for this study will be available at the European Nucleotide Archive ENA database (accession number PRJEB54646). The bionano raw data and assembled optical maps will be available at National Library for Biotechnology Information NCBI database (sample name PRJNA859008). All scripts and datasets will be available at Dryad database (doi:10.5061/dryad.pnvx0k6r0).

## Author contributions

L.F., L.A., G.S. & F.P.S. planned and designed the research. L.F. & L.A. conducted the fieldwork. L.F. coordinated the different collaborators involved in the project. L.F. improved the high-molecular weight genomic DNA extraction protocol for *de novo* sequencing and performed DNA extraction for pool-sequencing. L.P. preformed the Bionano optical mapping and scaffolding. W.Q. performed the bioinformatic analysis (assembly and annotation of the reference genome, Illumina read mapping and variant analysis), and wrote the methods related to sequencing and bioinformatics. L.F. performed the statistical analysis, the genome environmental association analysis, and the enrichment analysis. Q.R. performed *Brassica incana* - pollinator interaction analysis and wrote the related method part. L.F. wrote the manuscript, and all authors reviewed and edited the manuscript.

## Supplementary information

**Figure S1.**
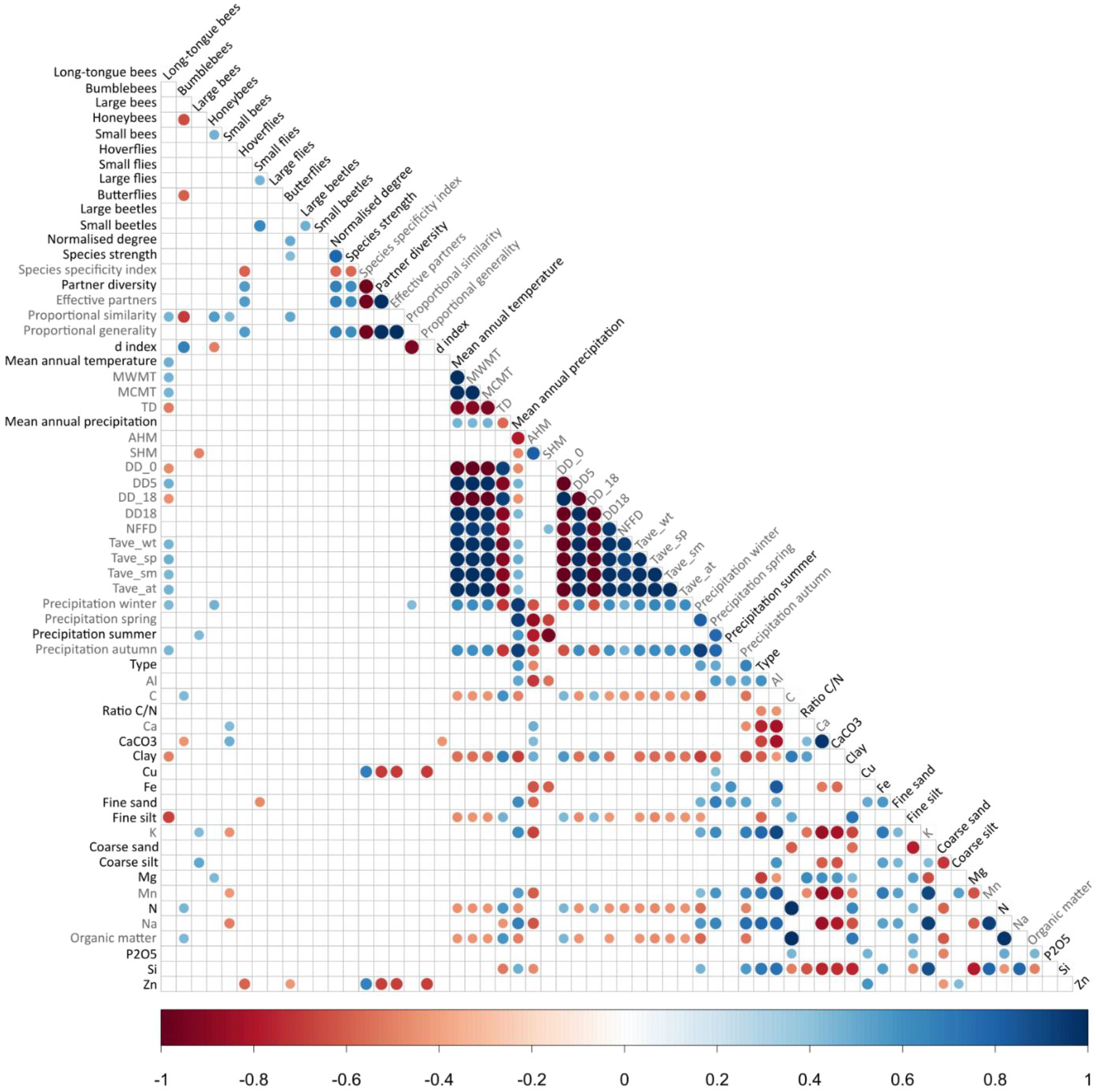
Matrix of spearman correlation on 61 ecological variables considered. Significant correlations are indicated by coloured dots. Not dots mean no significant correlations. The strength of the correlation is indicated by the size of the dots, and the direction by the blue and red gradient (gradient scale at the lower part of the figure). The traits indicated in grey were discarded from the genomic analysis due to high correlations with other traits (spearman *rho* > 0.8).

**Figure S2.**
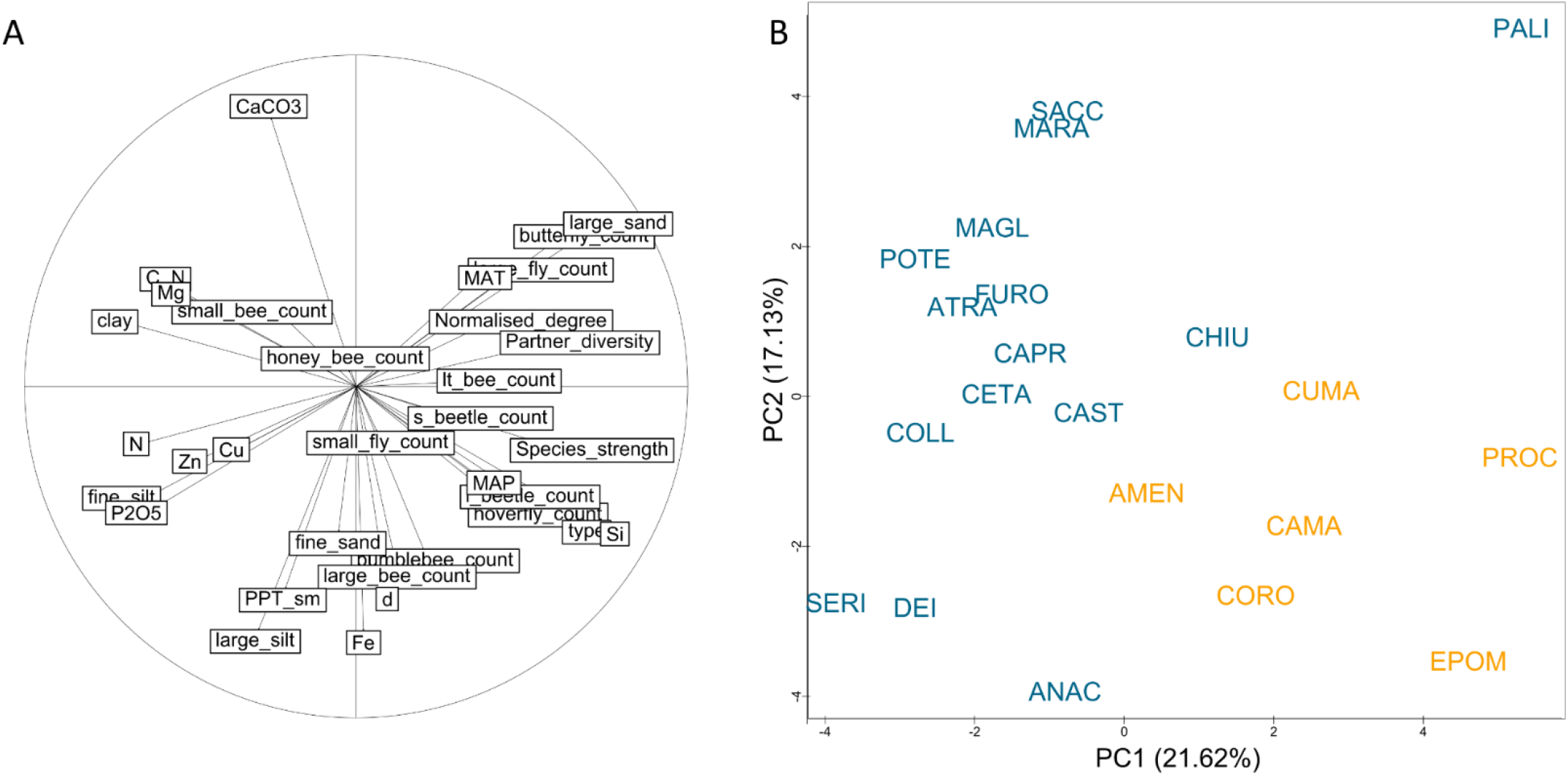
Ecological variation among 21 natural populations of *B. incana*. **(A)** Correlation plot from a principal component analysis preformed on 33 environmental variables with pairwise rho spearman < 0.8. Principal component PC1 and PC2 explained 21.62% and 17.13% respectively. **(B)** Position of the 21 natural populations of *B. incana* in ecological space. The populations in tuff soil are coloured in orange, and in limestone soil in blue.

**Figure S3.**
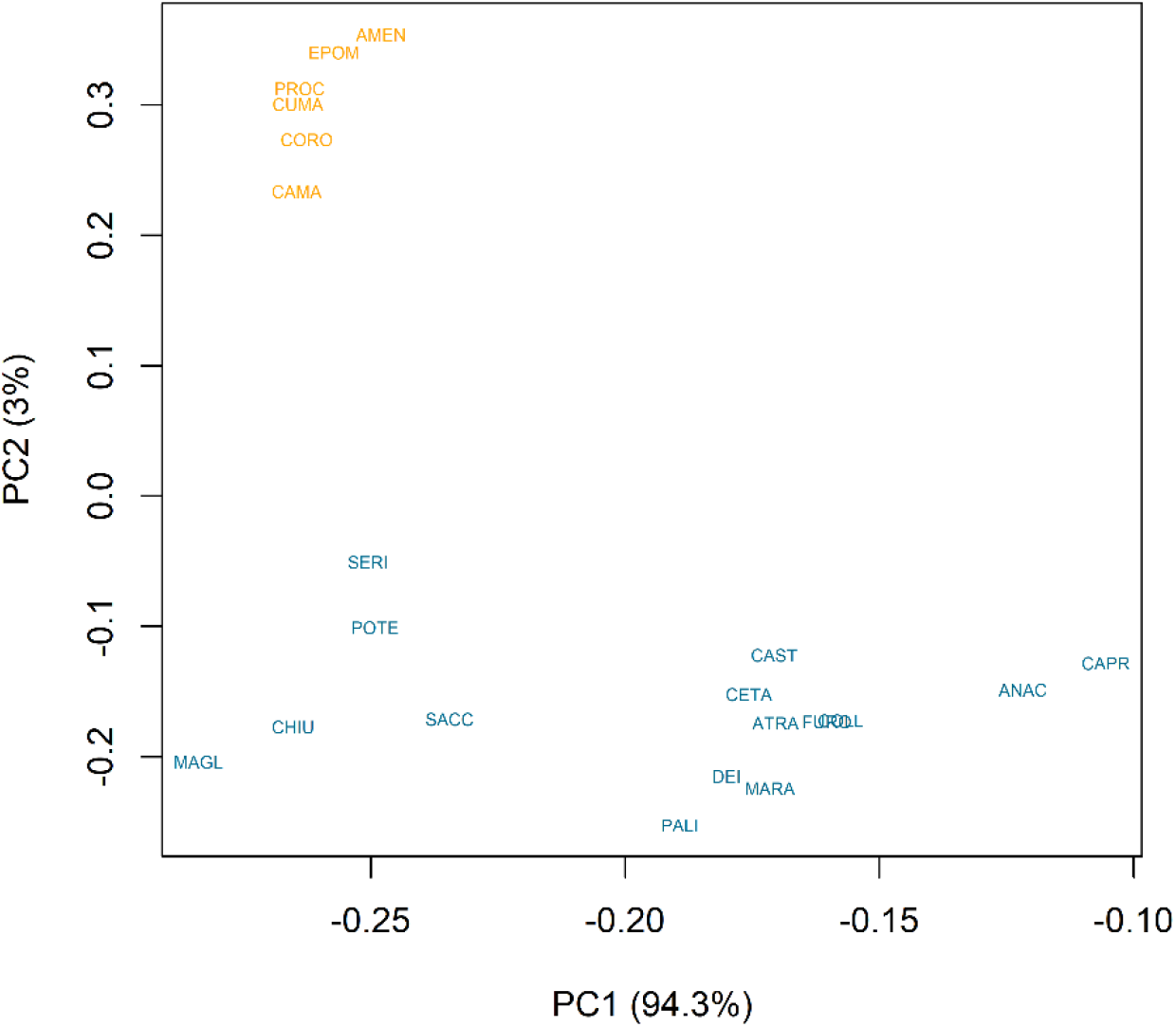
Position of 21 natural populations of *B. incana* in genomic space. Genomic variation was estimated using a singular value decomposition (SVD) of omega matrix using Baypass software from one sub-sample. The first PC_genomic_ explaining 94.3% of the genomic variance is represented on the *x-axis*, and the second PC_genomic_ on the *y-axis* explaining 3% of the genomic variance. The tuff and limestone soil of 21 populations are indicated in orange and in blue, respectively.

**Figure S4.**
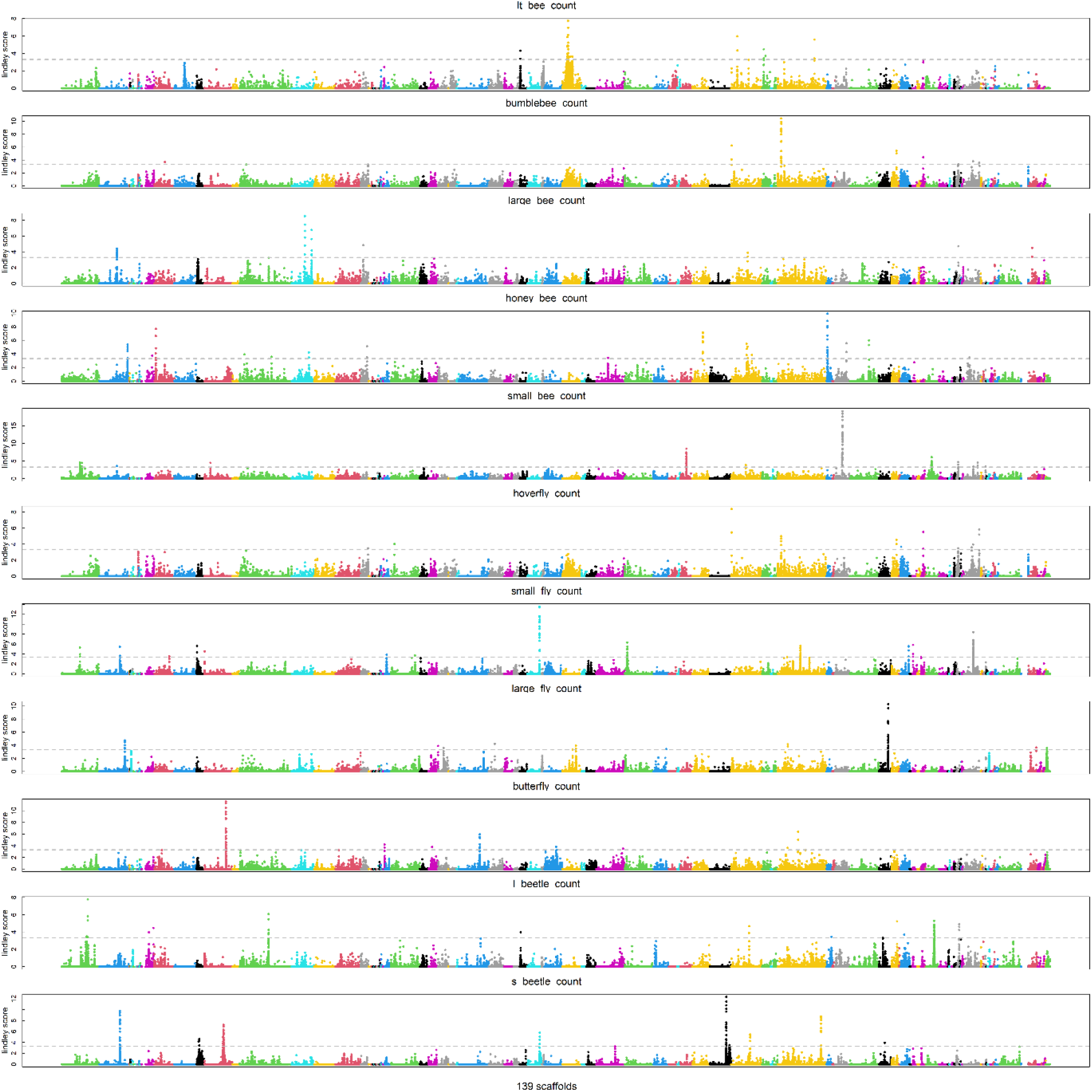
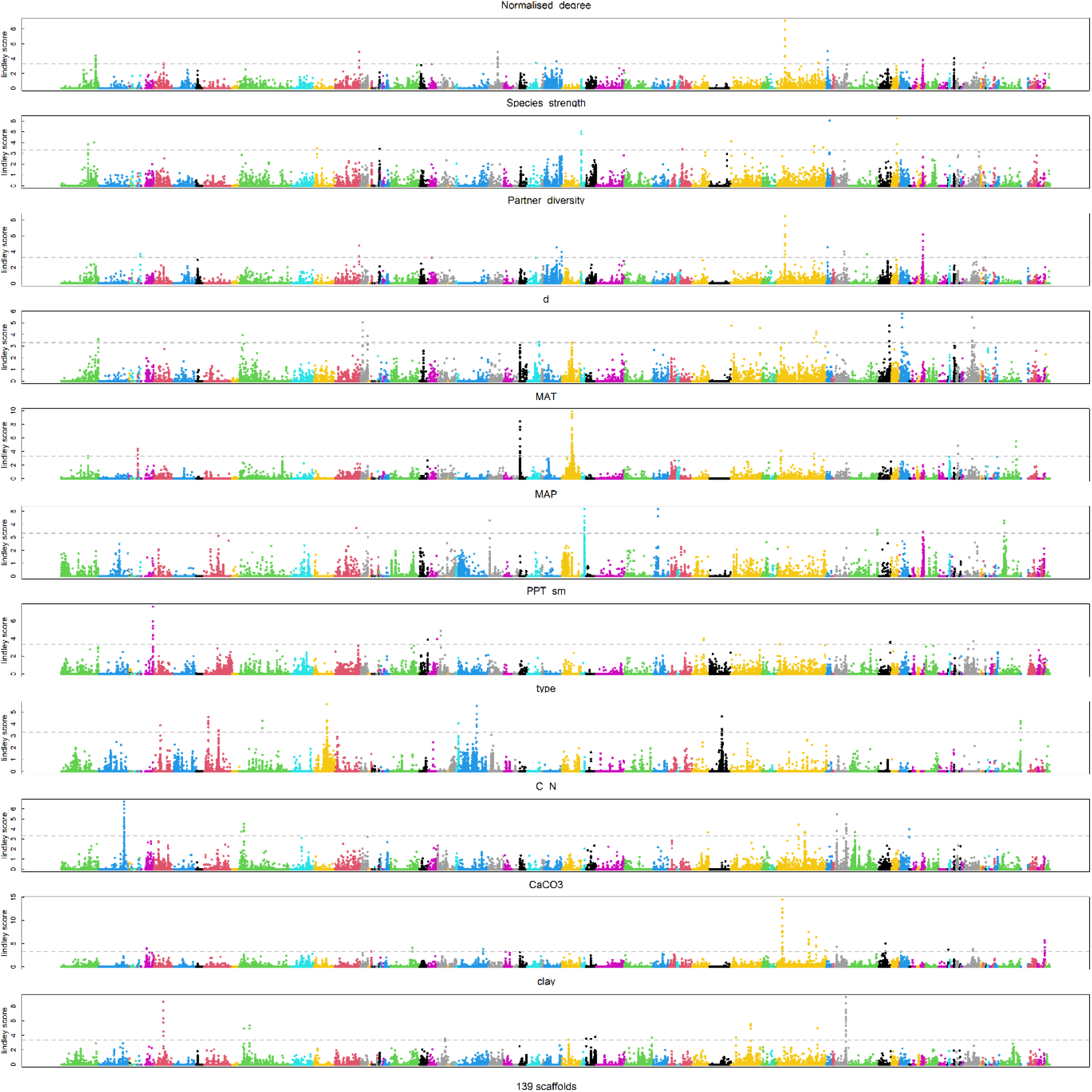
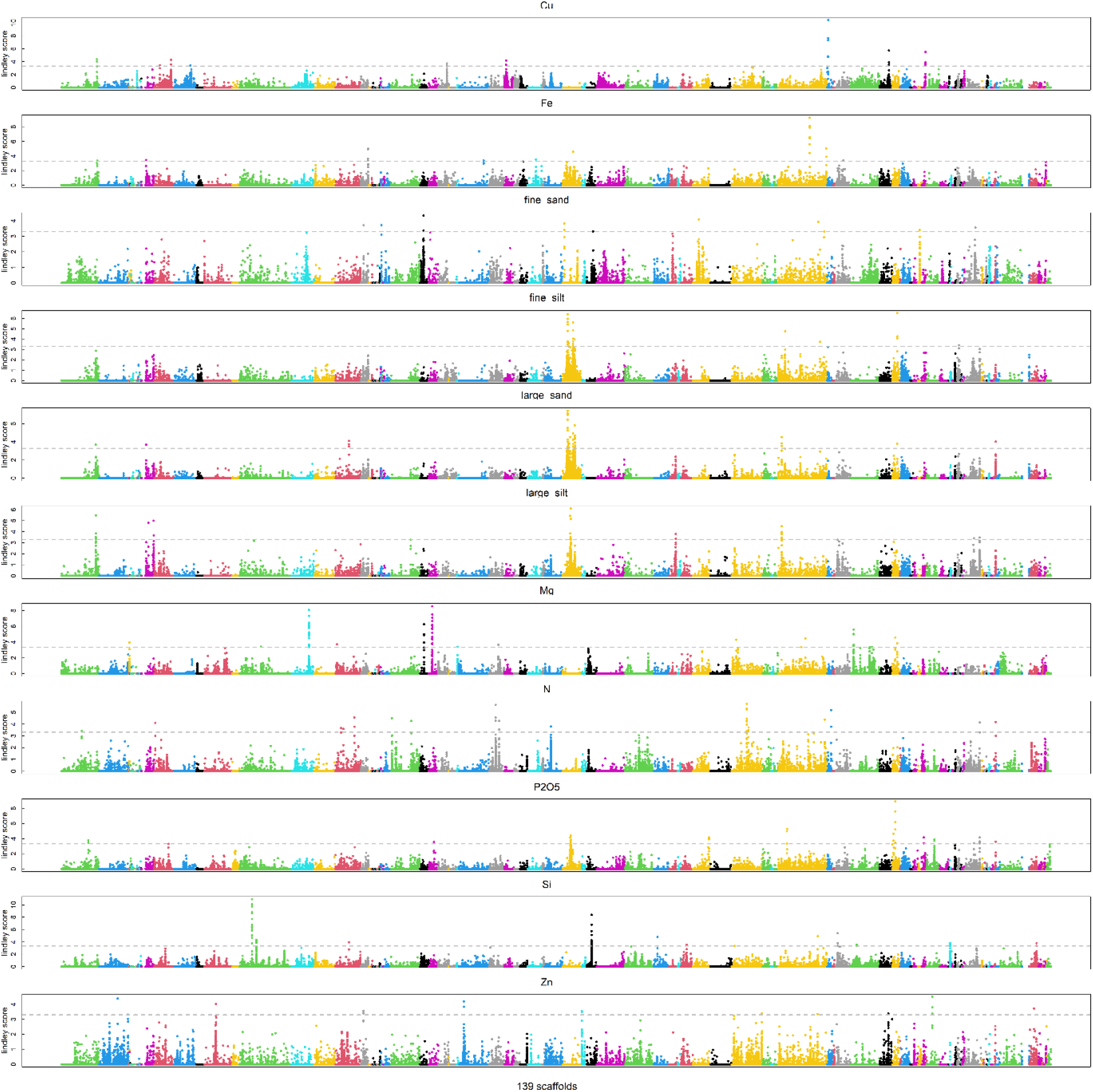
Manhattan plots of Genome-Environmental Association performed on 33 ecological variables. The x-axis represents the physical position of SNPs along the 139 super-scaffolds illustrated in colour. The *y-axis* is the Lindley score. The name of the ecological variable is indicated on the upper part of the Manhattan plot.

**Figure S5.**
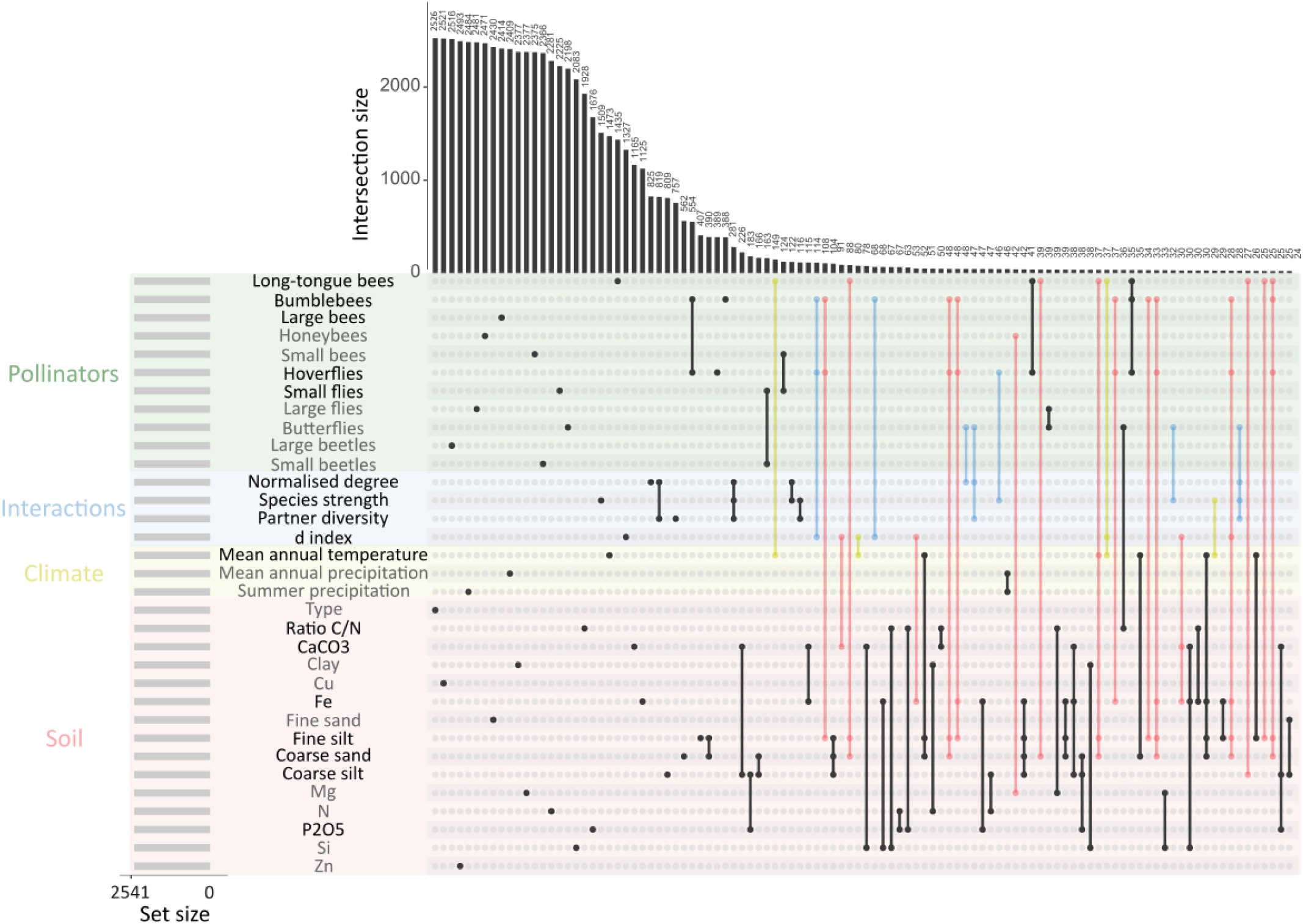
Illustration of the variability of SNPs involved in plant adaptation to a complex ecological network. The upset plot illustrates the specific SNPs to 33 ecological variables (only dots), and the shared SNPs among these ecological variables (dots linked with bar). The blue bars represent SNPs shared among *B. incana* responses to plant-pollinators interaction indices and categories of pollinators. The yellow bars represent the SNPs shared among the climate variables and pollinator community descriptors (categories and interactions). The red bars represent the SNPs shared among edaphic variables and the pollinator community descriptors. The top 0.05% SNPs of the highest association score were considered for 33 ecological variables listed in the left (*i*.*e*. set size = 2541 SNPs per ecological variable). Only the 156 first intercepts are shown (*i*.*e*. more than 1% of the set size). For instance, 1435 SNPs are unique to long-tongue bees, and 149 SNPs are shared between long-tongue bees and mean annual temperature. The ecological variables with non-significant enrichment in signature of selection have been shaded.

**Figure S6:**
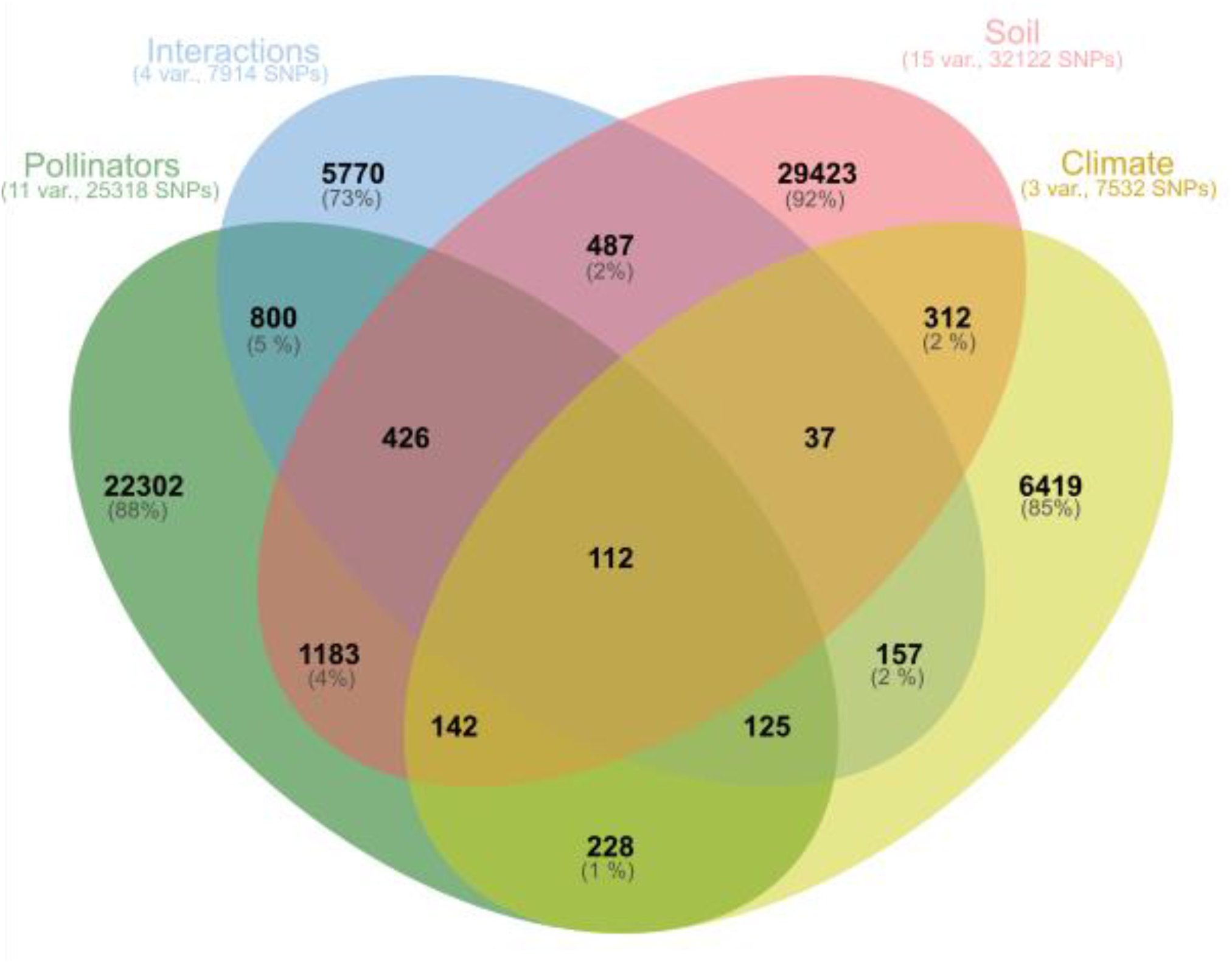
Illustration of the flexibility of genetic architecture in response to complex a ecological network. Venn diagram illustrating the shared top SNPs (0.05% of the highest local score) for the 33 ecological variables. The variables are grouped by main categories (pollinator categories in green, plant-pollinators interaction indices in blue, edaphic variables in red, and climatic variables in yellow). The number of variables and the total number of SNPs considered are indicated between parenthesis bellow each category of variables. The Venn diagram was draw using jvenn.toulouse.inra.fr website.

**Figure S7.**
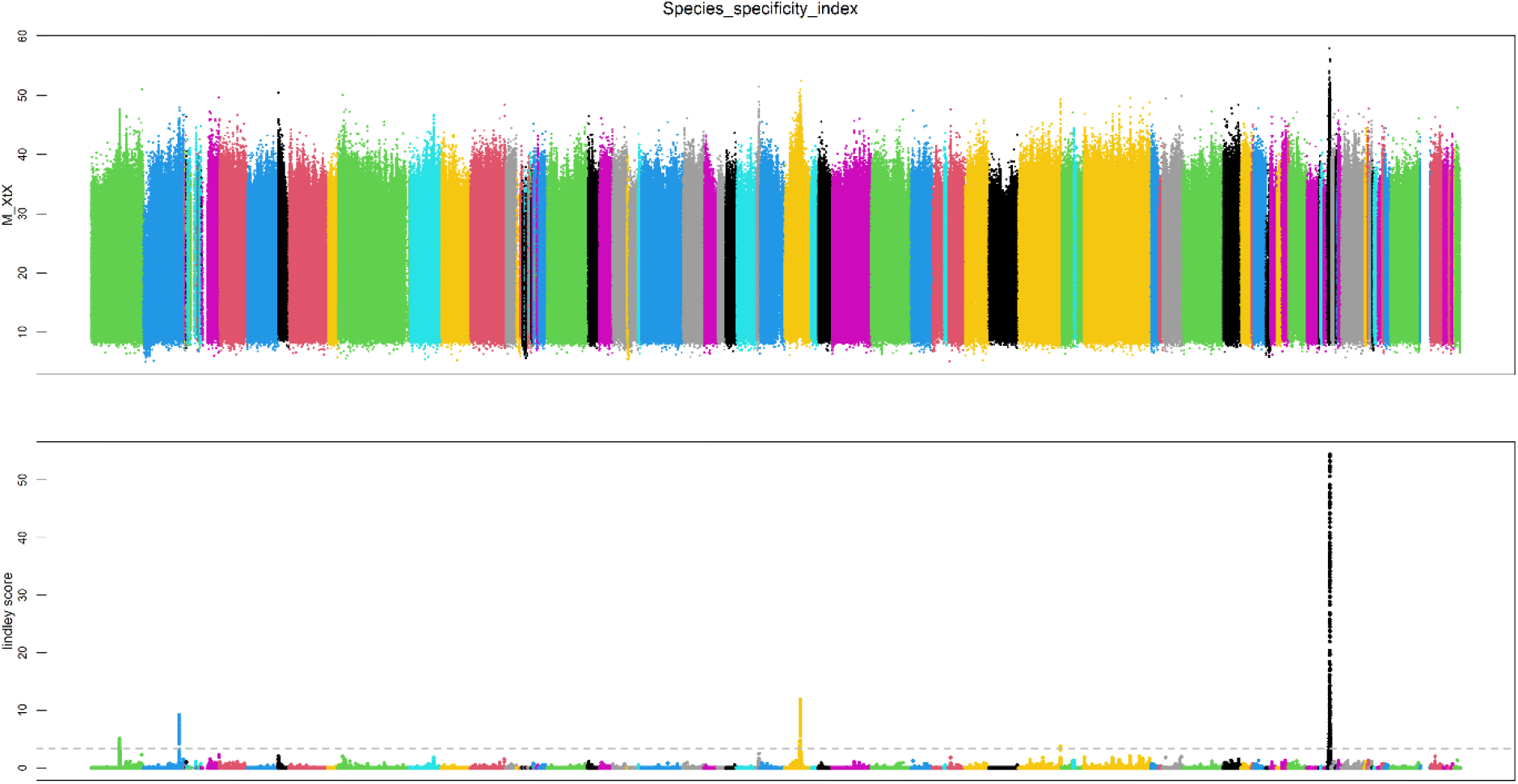
Manhattan plot of the index of genetic differentiation genetic (XtX). The upper panel illustrated the results obtained by Baybass analysis, and the lower panel those obtained by correcting with the local score method (*y-axis* is the Lindley score). The *x-axis* represents the physical regions of the SNPs along the 139 super-scaffolds.

**Table S1.** Description of 21 natural populations of *B. incana*.

| Pop. Name | Localities | latitude | longitude | Elevation | Substrate | Pop. Size 2018 |
| --- | --- | --- | --- | --- | --- | --- |
| PROC | Monte di Procida | 40.809370 | 14.044790 | 16 | Tuff | 20 |
| AMEN | Ischia, Lacco Ameno | 40.751632 | 13.896623 | 6 | Tuff | 30 |
| EPOM | Ischia, Vetta Epomeo | 40.730072 | 13.895328 | 767 | Tuff | 20 |
| CUMA | Mainland di Cuma | 40.850475 | 14.049944 | 41 | Tuff | 20 |
| CORO | Napoli, Coroglio-Nisida | 40.798367 | 14.175846 | 19 | Tuff | 40 |
| CAPR | Capri, Scala Fenicia | 40.556357 | 14.228237 | 180 | limestone | 100 |
| ANAC | Anacapri, Monte Solaro | 40.545771 | 14.223282 | 562 | limestone | 1000 |
| CHIU | Valico di Chiunzi | 40.719061 | 14.619117 | 648 | limestone | 15-30 |
| COLL | Colli-Positano | 40.619627 | 14.447449 | 253 | limestone | 15-30 |
| FURO | Furore | 40.614468 | 14.548246 | 91 | limestone | 15-2 |
| CAMA | Napoli, Camaldoli | 40,855181 | 14.207769 | 159 | Tuff | 20-30 |
| MAGL | Magliano Vetere | 40.342298 | 15.242826 | 622 | limestone | 15-20 |
| PALI | Palinuro, Arco naturale | 40.030830 | 15.308058 | 2 | limestone | 30-50 |
| ATRA | Atrani | 40.636737 | 14.610037 | 52 | limestone | 15 |
| SERI | Serino | 40.8204531 | 14.919611 | 708 | limestone | 10-15 |
| SACC | Sacco, Salerne | 40.386551 | 15.366420 | 505 | limestone | 50-100 |
| CAST | Castellammare di Stabia | 40.682200 | 14.439804 | 20 | limestone | 50 |
| DEI | Sentiero degli Dei | 40.625781 | 14.536430 | 649 | limestone | 15 |
| CETA | Cetara | 40.645236 | 14.699832 | 52 | limestone | 30 |
| POTE | Vietri di potenza | 40.570945 | 15.520259 | 359 | limestone | 30 |
| MARA | Maratea | 40.0421848 | 15.6525735 | 135 | limestone | 100 |

**Table S2.** Indices from the *Brassica incana* -- pollinator interaction analysis. The description of the indices is available in the methods and in Dormann 2011. In grey, the metrics discarded for the genomic analysis due to high correlation with other ecological factors (See Figure S1).

| Populations | Normalised degree | Species strength | Species specificity index | Partner diversity | Effective partners | Proportional similarity | Proportional generality | d |
| --- | --- | --- | --- | --- | --- | --- | --- | --- |
| AMEN | 0.42 | 0.18 | 0.73 | 0.85 | 2.35 | 0.43 | 0.38 | 0.21 |
| ANAC | 0.67 | 0.88 | 0.44 | 1.56 | 4.76 | 0.38 | 0.77 | 0.39 |
| ATRA | 0.67 | 0.44 | 0.58 | 1.23 | 3.41 | 0.66 | 0.55 | 0.13 |
| CAMA | 0.50 | 0.40 | 0.46 | 1.47 | 4.35 | 0.62 | 0.70 | 0.14 |
| CAPR | 0.25 | 0.09 | 0.79 | 0.63 | 1.88 | 0.44 | 0.30 | 0.20 |
| CAST | 0.58 | 0.27 | 0.53 | 1.34 | 3.80 | 0.76 | 0.61 | 0.07 |
| CETA | 0.75 | 0.68 | 0.66 | 1.19 | 3.28 | 0.51 | 0.53 | 0.20 |
| CHIU | 0.67 | 1.55 | 0.48 | 1.46 | 4.29 | 0.72 | 0.69 | 0.10 |
| COLL | 0.25 | 0.03 | 0.77 | 0.68 | 1.98 | 0.38 | 0.32 | 0.17 |
| CORO | 0.50 | 0.58 | 0.51 | 1.31 | 3.70 | 0.72 | 0.60 | 0.10 |
| CUMA | 0.58 | 0.56 | 0.54 | 1.32 | 3.74 | 0.74 | 0.60 | 0.09 |
| DEI | 0.17 | 0.04 | 0.72 | 0.64 | 1.89 | 0.14 | 0.31 | 0.32 |
| EPOM | 0.67 | 1.71 | 0.37 | 1.74 | 5.67 | 0.26 | 0.92 | 0.53 |
| FURO | 0.67 | 0.21 | 0.44 | 1.58 | 4.84 | 0.80 | 0.78 | 0.04 |
| MAGL | 0.67 | 0.67 | 0.47 | 1.48 | 4.41 | 0.78 | 0.71 | 0.06 |
| MARA | 0.50 | 0.42 | 0.53 | 1.26 | 3.52 | 0.65 | 0.57 | 0.12 |
| PALI | 0.67 | 0.67 | 0.42 | 1.61 | 5.02 | 0.66 | 0.81 | 0.14 |
| POTE | 0.50 | 0.10 | 0.46 | 1.43 | 4.19 | 0.69 | 0.68 | 0.09 |
| PROC | 0.83 | 1.49 | 0.39 | 1.80 | 6.06 | 0.70 | 0.98 | 0.09 |
| SACC | 0.67 | 0.48 | 0.55 | 1.33 | 3.77 | 0.70 | 0.61 | 0.11 |
| SERI | 0.58 | 0.58 | 0.69 | 1.08 | 2.95 | 0.48 | 0.48 | 0.23 |

**Table S3.** Sequence data collected for *de novo* genome assembly of *Brassica incana* from Pacbio and Illumina.

|  | PacBio CLR | Illumina PE reads |
| --- | --- | --- |
| Number of reads | 2,481,304 | 249,786,052 |
| Number of bases (bp) | 47,372,198,992 | 74,935,815,600 |
| Read N50 | 17 kbp | 2X150 bp |
| Estimated coverage* | 73 X | 115 X |

\*assumed genome size of 650 Mbp

**Table S4.** Bionano row molecule data collected for hybrid scaffolding of *Brassica incana* contigs.

| Protocol |  | NLRS | DLS |
| --- | --- | --- | --- |
| Enzyme |  | Nb.BspQI | DLE-1 |
| Molecule >= 20 kbp | Total length (Mbp) | 1,694 | 2,278 |
|  | N50 (Mbp) | 0.101 | 0.123 |
| Molecule >= 150 kbp | Total length (Mbp) | 549 | 972 |
|  | N50 (Mbp) | 233 | 0.267 |
|  | Label density | 5.43/100 kbp | 5.03/100 kbp |
| Effective coverage |  | 340.88 | 34.82 |

**Table S5.**
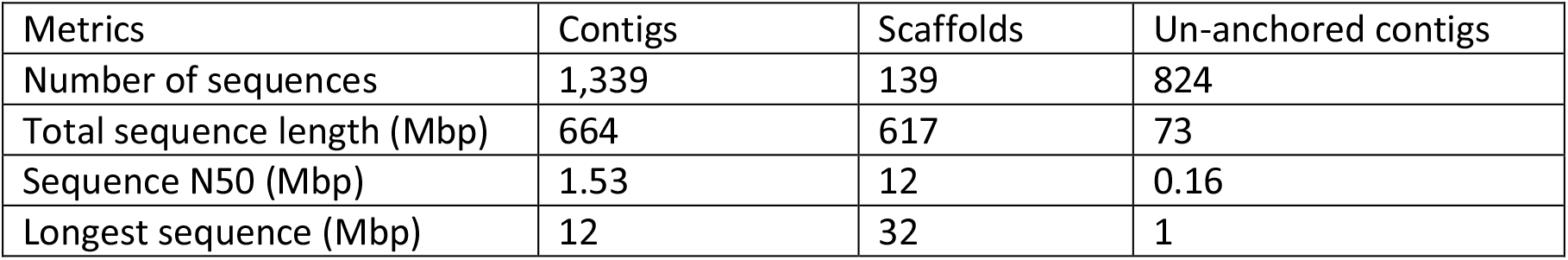
Final assembly statistics of *Brassica incana* contigs and scaffolds.

**Table S6.**
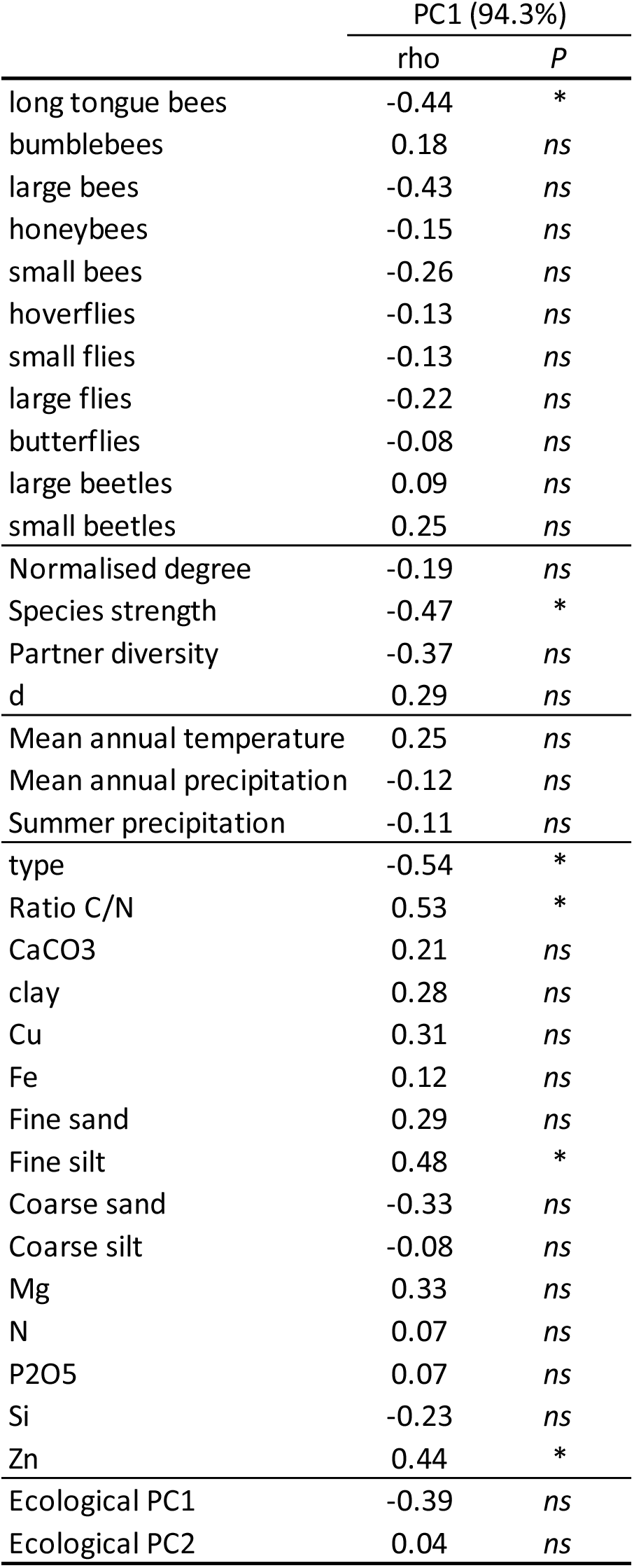
Spearman correlation between genomic variance (SVG) and 33 ecological variables and PC1 and PC2 from the principal analysis performed on 33 ecological variables.

|  | PC1 (94.3%) |  |
| --- | --- | --- |
|  | rho | P |
| long tongue bees | -0.44 | * |
| bumblebees | 0.18 | ns |
| large bees | -0.43 | ns |
| honeybees | -0.15 | ns |
| small bees | -0.26 | ns |
| hoverflies | -0.13 | ns |
| small flies | -0.13 | ns |
| large flies | -0.22 | ns |
| butterflies | -0.08 | ns |
| large beetles | 0.09 | ns |
| small beetles | 0.25 | ns |
| Normalised degree | -0.19 | ns |
| Species strength | -0.47 | * |
| Partner diversity | -0.37 | ns |
| d | 0.29 | ns |
| Mean annual temperature | 0.25 | ns |
| Mean annual precipitation | -0.12 | ns |
| Summer precipitation | -0.11 | ns |
| type | -0.54 | * |
| Ratio C/N | 0.53 | * |
| CaCO3 | 0.21 | ns |
| clay | 0.28 | ns |
| Cu | 0.31 | ns |
| Fe | 0.12 | ns |
| Fine sand | 0.29 | ns |
| Fine silt | 0.48 | * |
| Coarse sand | -0.33 | ns |
| Coarse silt | -0.08 | ns |
| Mg | 0.33 | ns |
| N | 0.07 | ns |
| P2O5 | 0.07 | ns |
| Si | -0.23 | ns |
| Zn | 0.44 | * |
| Ecological PC1 | -0.39 | ns |
| Ecological PC2 | 0.04 | ns |

**Table S7.** Enrichment in signature of selection for 33 ecological variables including pollinator categories (11 variables), plant-pollinators interaction indices (4 variables), climatic (3 variables) and edaphic variables (15 variables) in the 0.05% upper tail of the Lindley score distribution in the 0.05% upper tail of the genome-wide spatial differentiation (XtX) distribution.

| Traits | ntops | Enrichment | pvalue |
| --- | --- | --- | --- |
| Long-tongue bees | 21 | 16.53 | ** |
| Bumblebees | 23 | 18.11 | *** |
| Large bees | 15 | 11.81 | ** |
| Honeybees | 1 | 0.79 | *ns* |
| Small bees | 1 | 0.79 | *ns* |
| Hoverflies | 25 | 19.68 | *** |
| Small flies | 6 | 4.72 | * |
| Large flies | 0 | 0.00 | *ns* |
| Butterflies | 2 | 1.57 | *ns* |
| Large beetles | 0 | 0.00 | *ns* |
| Small beetles | 0 | 0.00 | *ns* |
| Normalised degree | 19 | 14.96 | ** |
| Species strength | 6 | 4.72 | * |
| Partner diversity | 20 | 15.74 | *** |
| d index | 31 | 24.40 | *** |
| Mean annual temperature | 109 | 85.81 | *** |
| Mean annual precipitation | 4 | 3.15 | *ns* |
| Summer precipitation | 0 | 0.00 | *ns* |
| type | 0 | 0.00 | *ns* |
| Ratio C/N | 13 | 10.23 | ** |
| CaCO3 | 12 | 9.45 | ** |
| clay | 0 | 0.00 | *ns* |
| Cu | 0 | 0.00 | *ns* |
| Fe | 39 | 30.70 | *** |
| Fine sand | 3 | 2.36 | *ns* |
| Fine silt | 59 | 46.45 | *** |
| Coarse sand | 62 | 48.81 | ** |
| Coarse silt | 31 | 24.40 | ** |
| Mg | 1 | 0.79 | *ns* |
| N | 3 | 2.36 | *ns* |
| P2O5 | 15 | 11.81 | ** |
| Si | 0 | 0.00 | *ns* |
| Zn | 3 | 2.36 | *ns* |

**Table S8.** Candidate genes is available in a separated file.

